# Behavioral dysfunction but not wasting in cancer cachexia is driven by altered tryptophan metabolism

**DOI:** 10.64898/2026.10.08.757616

**Authors:** Damon Fard, Edisa Pirani, Maria Gomez-Jenkins, Sereno Lopez-Darwin, Maria Ibrahim, Lilia Schell, Charlotte Larkin, Clara Gorgui, Clara Berdasco, Zhixian Hu, Trishna Das, Jesse Powers, Tracy G. Anthony, Joshua D. Rabinowitz, Benjamin J Raphael, Jessie Yanxiang Guo, Marcus D. Goncalves, Tobias Janowitz, Eileen White

## Abstract

Cancer cachexia causes pathological weight loss, anorexia, and depressive symptoms, culminating in increased mortality. Using mouse models of pancreatic cancer cachexia, we identify tumor and hepatic inflammatory signaling, multi-organ energy charge depletion, and altered tryptophan metabolism. Metabolomic profiling and stable isotope tracing revealed altered tryptophan fates: serotonin depletion and kynurenine accumulation across tumor, liver, muscle and spleen. Kynurenine accumulation involved accelerated conversion of tryptophan to kynurenine with slowed downstream conversion to kynurenic acid. Restoring serotonin activity pharmacologically or metabolically normalized feeding, locomotion, and depression-like behavior, but wasting, tumor burden, and survival remained unchanged. Blocking kynurenine production or kynurenine-mediated AhR activation pharmacologically produced similar results. Thus, in cancer cachexia, the tryptophan pathway is subverted to deplete serotonin and accumulate kynurenine. This subversion contributes to cachexia-associated behavioral and feeding defects but not lethal systemic catabolism.

**Highlights:**

- Tumors divert tryptophan from serotonin toward kynurenine accumulation in cachexia
- Serotonin depletion contributes to behavioral alterations in cancer cachexia
- Kynurenine accumulation reduces feeding and activity via aryl hydrocarbon receptor
- Combined pathway targeting preserves fat mass but wasting remains unaltered

**Graphical Abstract:** 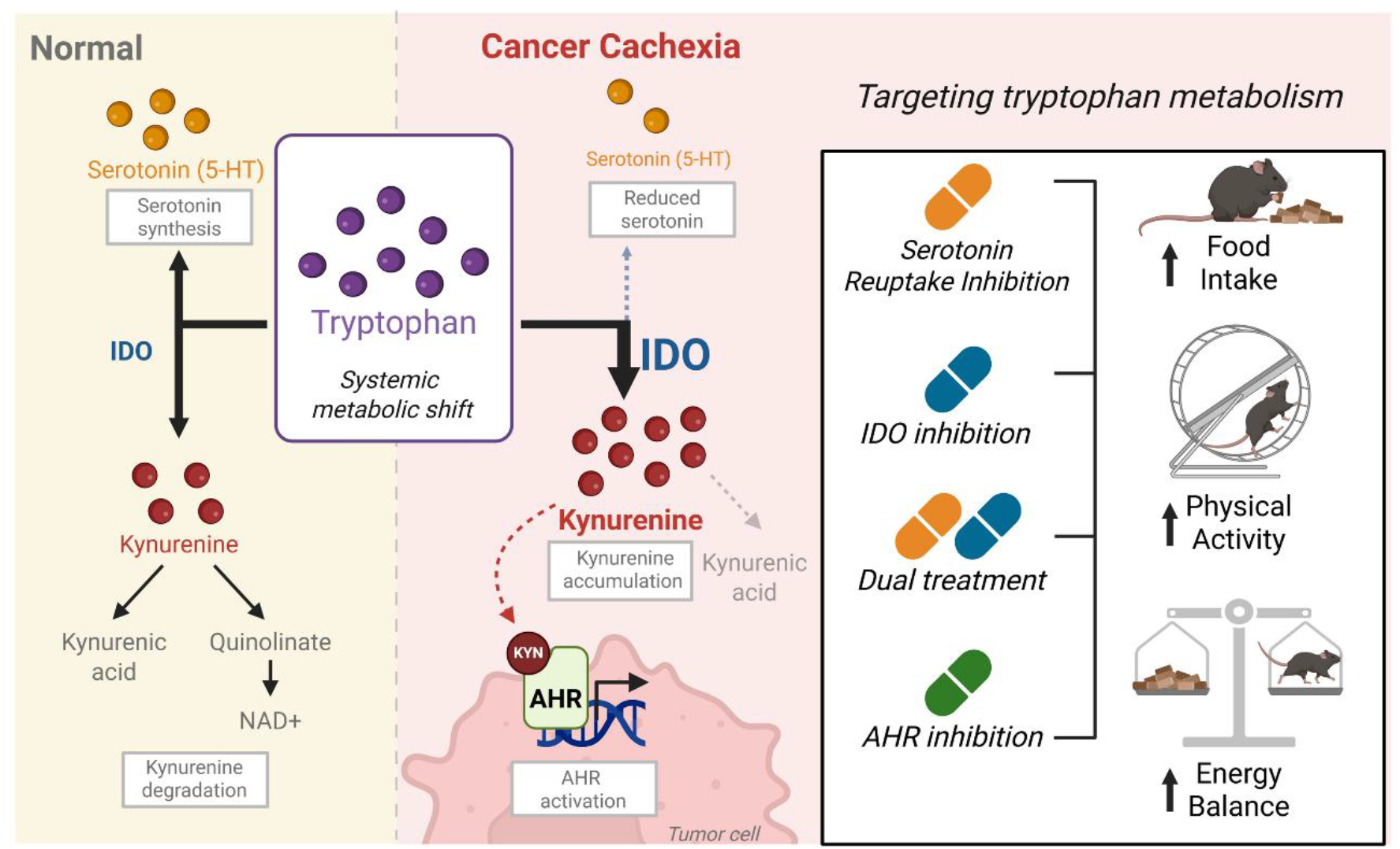

## Introduction

Cancer cachexia is a debilitating syndrome of severe muscle and fat loss, systemic inflammation, metabolic dysfunction, depression, anorexia-associated reduced calorie intake, and reduced activity, affecting up to 80% of pancreatic cancer patients^1–5^. Clinically, it is linked to poorer treatment outcomes, reduced quality of life, elevated mortality risk, and activity levels similar to those with spinal cord injury or cerebral palsy^6,7^. Despite this, the metabolic and molecular mechanisms underlying its diverse manifestations remain poorly understood, and effective therapies are lacking.

Cachexia is a systemic condition involving coordinated interactions among multiple tissues. Cancer alters the function of metabolic organs including liver, skeletal muscle, and adipose tissue, indicating that the syndrome arises from tumor-induced cross-talk between organs rather than changes in a single tissue. Body wasting is thought to be driven by tumor-derived signals that disrupt host metabolic homeostasis. Tumor secreted cytokines, including interleukin-6 (IL-6)^8,9^, Tumor necrosis factor-alpha (TNF-α)^10,11^, and leukemia inhibitory factor (LIF)^12,13^ can alter metabolic programs in distant organs such as skeletal muscle, adipose tissue, and liver. Growth differentiation factor 15 (GDF15), a stress-induced cytokine, has emerged as a key regulator of cachexia-associated anorexia through activation of the GDF15–GFRAL axis^14^. Blocking this pathway potentially improves appetite, body weight, and cachexia related symptoms in preclinical and clinical settings where GDF-15 is elevated^15^. However, these effects are incomplete and do not reverse cachexia progression. Tumors thereby establish a feed forward metabolic cycle promoting progressive tissue catabolism and body weight loss. Circulating metabolites, including amino acids^16,17^, products of amino acid metabolism^18,19^, and lipid species^20^, are altered in cachectic patients and distinguish stable weight from cancer-associated wasting, suggesting use as biomarkers of disease progression.

Cancer cachexia also includes prominent sickness behavior that directly worsens wasting. Patients frequently develop anorexia, fatigue, reduced motivation, depression-like symptoms, and anhedonia, indicating that cachexia affects both peripheral metabolism and central behavioral regulation^4,21–24^. These alterations are clinically important, reduced appetite accelerates negative energy balance and reduced activity impairs functional status, reinforcing systemic wasting.

Most studies have focused on skeletal muscle and adipose wasting, while systemic metabolomic alterations that drive cachexia remained poorly defined^25–27^. To define systemic metabolic alterations across tissues during disease progression in cancer cachexia, we integrated multi-tissue metabolomics and gene expression with cytokine and chemokine profiling in genetically engineered and orthotopic pancreatic cancer mouse models. Together, these findings define a conserved inflammatory metabolic program in pancreatic cancer cachexia and reveal that altered tryptophan metabolism drives behavioral symptoms mechanistically separable from lethal metabolic decline.

## Results

### Pancreatic cancer drives progressive tissue wasting and disrupts whole-body energy balance

In this study, we used two mouse models of pancreas ductal adenocarcinoma (PDAC), GEM model KPC (KPC LSL-Kras^G12D/+^; LSL-Trp53^R172H/+^; Pdx1-Cre) and an orthotopic model generated using two independent KPC-derived mouse pancreatic adenocarcinoma cell lines implanted in the pancreas of wild-type C57BL/6J mice. Mice were followed longitudinally by EchoMRI, ultrasound, and metabolic-cage analysis; tumor bearing mice with 5%–15% body weight loss were classified as pre-cachectic and those with >15% loss as severely cachectic (Figure 1A). In both models, body weight, lean mass, and especially fat mass were significantly reduced at the cachectic endpoint relative to controls (Figures 1B and 1C), showing that spontaneous and orthotopic PDAC produce a common wasting phenotype with pronounced fat depletion.

**Figure 1.**
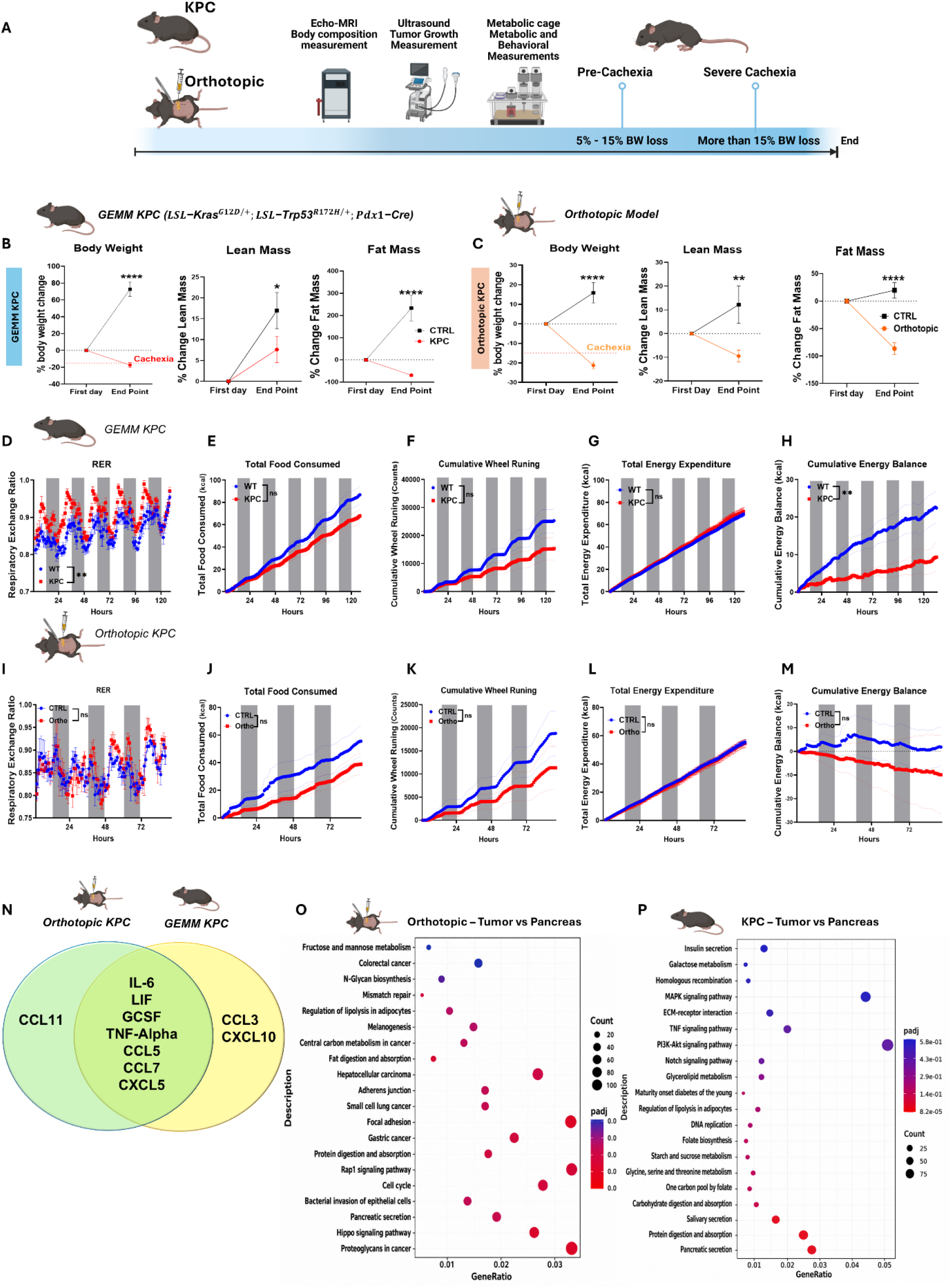
Systemic and metabolic features of PDAC cachexia. (A) GEMM KPC and orthotopic pancreatic ductal adenocarcinoma (PDAC) models and timeline. PreCCX, pre-cachexia (5%–15% body weight loss); SCCX, severe cachexia (>15%). (B and C) Change in body weight, lean mass, and fat mass from baseline to endpoint in control versus GEMM KPC (B) or orthotopic PDAC (C) mice. *n* = 5 (B) and 4 (C) mice per group. (D–H) Metabolic-cage analysis of mixed-sex control versus GEMM KPC mice: respiratory exchange ratio (RER; D), food intake (E), wheel running (F), energy expenditure (G), and energy balance (H). *n* = 8 mice. (I–M) Same for orthotopic PDAC-bearing mice: RER (I), food intake (J), wheel running (K), energy expenditure (L), and energy balance (M). *n* = 4 mice. (N) Serum cytokines significantly altered in orthotopic PDAC and/or GEMM KPC mice versus controls. *n* = 15 KPC, 16 orthotopic, and 6 control mice per model. (O and P) Pathway enrichment of differentially expressed genes in orthotopic (O) and GEMM KPC (P) tumors versus normal pancreas. *n* = 3 mice per group. Data are mean ± SEM; two-way repeated-measures (RM) ANOVA with uncorrected Fisher’s LSD test (B and C), two-way RM ANOVA/mixed-effects model with Geisser–Greenhouse correction and Šídák’s test (D–M), or unpaired Student’s *t* test (N). \**p* < 0.05, \*\**p* < 0.01, \*\*\**p* < 0.001, \*\*\*\**p* < 0.0001. See also Figure S1.

To quantify systemic metabolic alterations during cachexia, we monitored mice in Promethion metabolic cages for substrate utilization, energy expenditure, food intake, and activity. Respiratory exchange ratio (RER) reflects fuel preference, with values near 1.0 indicating greater carbohydrate oxidation and values near 0.7 greater lipid oxidation. In both models, tumor bearing mice showed a concordant metabolic and behavioral phenotype that was evident over the short monitoring period and became more pronounced with disease progression. In KPC mice, RER was elevated, while cumulative food intake and voluntary wheel running progressively diverged from wild-type (WT) controls (Figures 1D–1F). Despite these reductions, total energy expenditure was largely unchanged (Figure 1G), resulting in a progressively lower cumulative energy balance (Figure 1H). The orthotopic model recapitulated this pattern, RER differed only modestly (Figure 1I), whereas cumulative food intake, wheel running progressively declined relative to controls (Figures 1J and 1K). Energy expenditure remained comparable to controls (Figure 1L), whereas cumulative energy balance became progressively more negative (Figure 1M). Thus, both models converge on a similar progressive phenotype, with declining food intake and spontaneous activity as shared features of disease progression.

### Conserved inflammatory signature across pancreatic cancer cachexia models

Cancer cachexia is characterized by chronic, systemic inflammation with dysregulated cytokine and chemokine signaling^28,29^, which drives tumor–host communication and contributes to metabolic dysfunction and tissue catabolism during disease progression^8,17^. To define the inflammatory landscape, we profiled circulating cytokines and chemokines by multiplex Luminex across defined stages of cachexia. Eight analytes were significantly altered in the orthotopic model and nine in the KPC model (Figure S1A and S1B). Seven were shared, indicating a conserved inflammatory signature of IL-6, LIF, G-CSF, TNF-α, CCL5, CCL7, and CXCL5, whereas CCL11 was orthotopic-specific and CCL3 and CXCL10 were KPC-specific (Figure 1N). IL-6 and LIF regulate inflammatory signaling and metabolic reprogramming linked to muscle proteolysis, adipose remodeling, and hepatic acute-phase responses^13,30^. TNF-α or cachectin promotes anorexia, systemic inflammation, and NF-κB-driven proteolysis^31,32^. CCL5 mediates tumor–immune crosstalk promoting catabolic signaling^33^, and CCL7 recruits macrophages, monocytes, and lymphocytes to remodel the tumor microenvironment^34^. Although each mediator has been studied individually in cachexia, our data show that they converge into a shared circulating inflammatory signature across two distinct models.

To define tumor transcriptional changes, we performed bulk RNA sequencing of tumor and matched pancreas from both models with pathway enrichment analysis. Orthotopic tumors showed enrichment of central carbon metabolism, N-glycan biosynthesis, cell-cycle programs, nutrient digestion and absorption, and multiple cancer-associated pathways versus pancreas (Figure 1O). KPC tumors showed similarly broad but distinct remodeling, including MAPK, PI3K-AKT, TNF, and Notch signaling with multiple metabolic pathways (Figure 1P). Despite differences in tumor origin and architecture, both models converged on coordinated metabolic and inflammatory transcriptional programs within the tumor. With the systemic cytokine and chemokine changes, these findings pointed to a broader tumor host response, prompting us to examine host metabolic tissues.

### Pancreatic cancer cachexia engages a multi-organ inflammatory program that extends to hepatocytes

We next asked whether tumor intrinsic pathway remodeling (Figures 1O and 1P) extended to distal organs. Bulk RNA-seq of spleen, inguinal white adipose tissue (iWAT), kidney, and quadriceps from KPC and orthotopic mice revealed inflammatory or stress transcriptional programs across all peripheral tissues examined (Figures S2A-S2D). Cytokine–cytokine receptor interaction was enriched in spleen, iWAT, and kidney in both models, whereas quadriceps showed enrichment of autophagy, mitophagy, and proteasome pathways, with induction of atrophy genes *Trim63*, *Fbxo32*, *Foxo3*, and *Mstn* (Figures S2E and S2F). Tumors also lost exocrine identity, suppressing digestive enzyme genes (Figure S2G), relevant given that exocrine pancreatic dysfunction impairs nutrient digestion and contributes to peripheral tissue wasting^35^.

In liver, immune pathways were most prominent in KPC mice, including cytokine–cytokine receptor, chemokine, and JAK–STAT signaling, while both models shared enrichment of hepatocyte-associated metabolic pathways, including cytochrome P450-dependent xenobiotic and drug metabolism, retinol metabolism, and steroid hormone biosynthesis (Figures 2A and 2B). Acute-phase genes, including *Lcn2* and *Orm1* in both models and fibrinogen genes (*Fga*, *Fgb*, *Fgg*) in orthotopic mice, were induced relative to controls, although induction varied between animals (Figures 2C and 2D). This mirrors our Luminex data showing elevated circulating IL-6 and TNF-α, placing the liver as a key downstream target of the systemic cytokine surge^36,37^, consistent with prior reports of widespread cachexia associated inflammation with a central hepatic role^38^.

**Figure 2.**
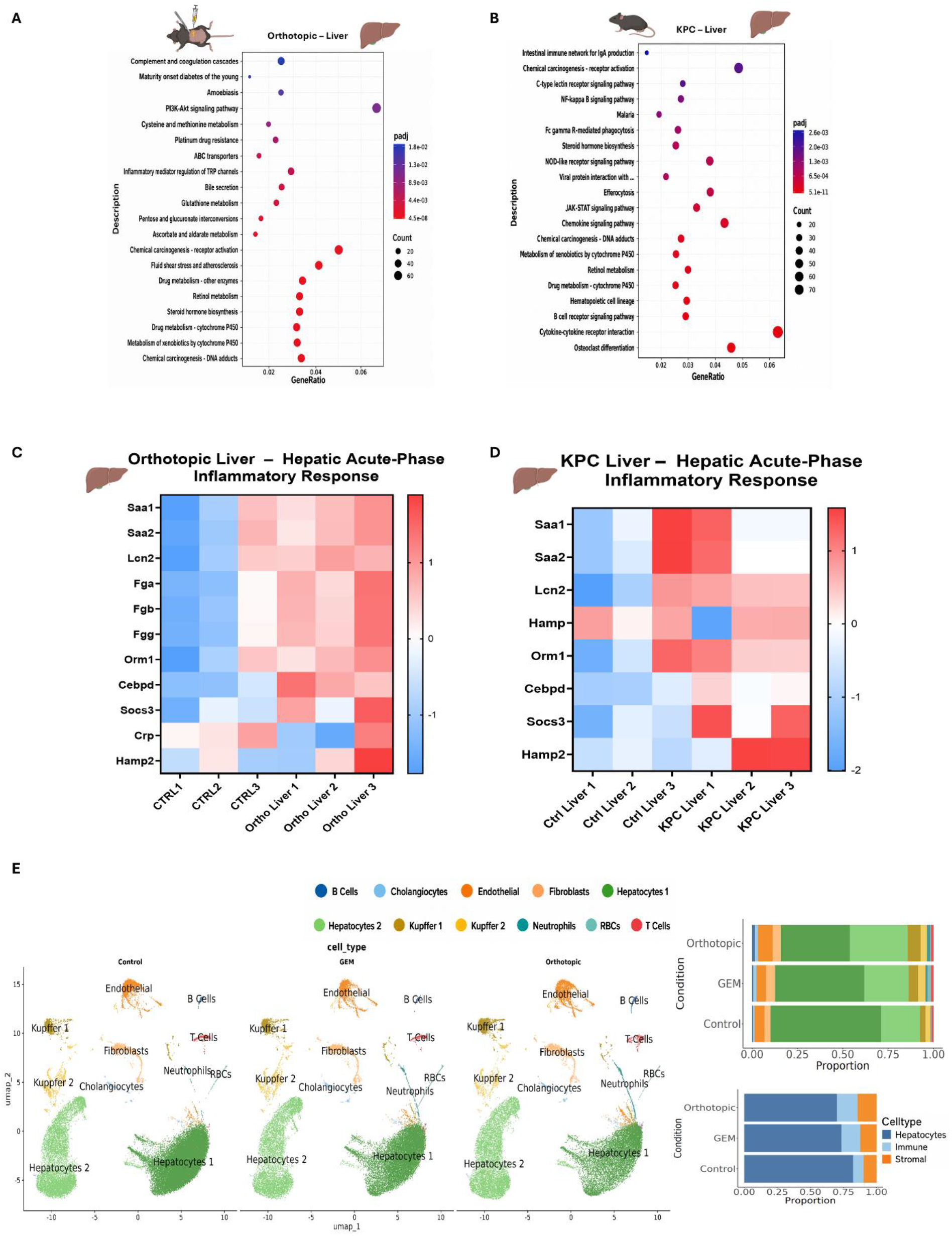
Hepatic inflammatory remodeling in PDAC cachexia. (A and B) Pathway enrichment of differentially expressed liver genes in orthotopic PDAC (A) and GEMM KPC (B) mice versus controls. (C and D) Hepatic acute-phase and inflammatory-response genes in orthotopic PDAC (C) and GEMM KPC (D) mice versus controls. (E) Liver single-nucleus RNA-seq of control, GEMM KPC, and orthotopic PDAC mice: UMAPs and cell-type and compartment proportions. *n* = 3 per group (A–E). See also Figure S2.

To resolve whether this response was tissue-wide or cell specific, we performed single-nucleus RNA-seq of livers from control, KPC, and orthotopic mice. Tumor bearing livers showed reduced hepatocyte fractions and expanded immune populations (Figure 2E), consistent with inflammation-associated hepatic remodeling^39–41^. Hepatocytes 1 decreased while Hepatocytes 2 relatively expanded in both models, most prominently in orthotopic livers (Figure 2E). Gene Ontology analysis showed acute-phase inflammatory enrichment in Hepatocytes 1, whereas Hepatocytes 2 instead showed metabolic programs, including amino acid catabolism and small-molecule metabolism (Figure S2H). Thus, the hepatic response extends beyond systemic inflammation to distinct hepatocyte subpopulations. We next performed multi-tissue metabolomic profiling to test for accompanying metabolic alterations.

### Tumor-driven alteration of tryptophan metabolism toward kynurenine defines a shared systemic signature of pancreatic cancer cachexia

We next asked whether this remodeling was accompanied by coordinated metabolic alterations. Tryptophan sits at a major metabolic branchpoint, supporting serotonin biosynthesis, kynurenine metabolism and downstream NAD+ production, and microbiota-derived indole pathways (Figure 3A). We therefore performed metabolomic profiling of serum and multiple tissues in KPC and orthotopic models across progression from pre-cachectic to severely cachectic states (Figures 3B–3I; Figures S3–S6). Across all tissues, cachexia converged on coordinated diversion of tryptophan away from serotonin synthesis and toward kynurenine. In serum, serotonin declined markedly while kynurenine rose progressively with severity, following nearly identical trajectories in both models (Figures 3B and 3F). Kynurenine also accumulated in tumor (Figures 3C and 3G), liver (Figures 3D and 3H), and spleen (Figures 3E and 3I), making it the most reproducible metabolic alteration observed. Despite this, downstream kynurenic acid, quinolinate, and indolelactic acid remained unchanged, revealing a metabolic bottleneck after kynurenine. The broader serum metabolome changed progressively with disease severity, with pre-cachectic mice occupying an intermediate state. Orthotopic PDAC mice also showed reduced succinate, fumarate, and α-ketoglutarate and increased 3-hydroxybutyrate, whereas KPC mice showed reduced α-ketoglutarate and malate and increased 2-hydroxybutyrate, accompanied by global separation by PCA (Figures S3A-S3B and S5A-S5B).

**Figure 3.**
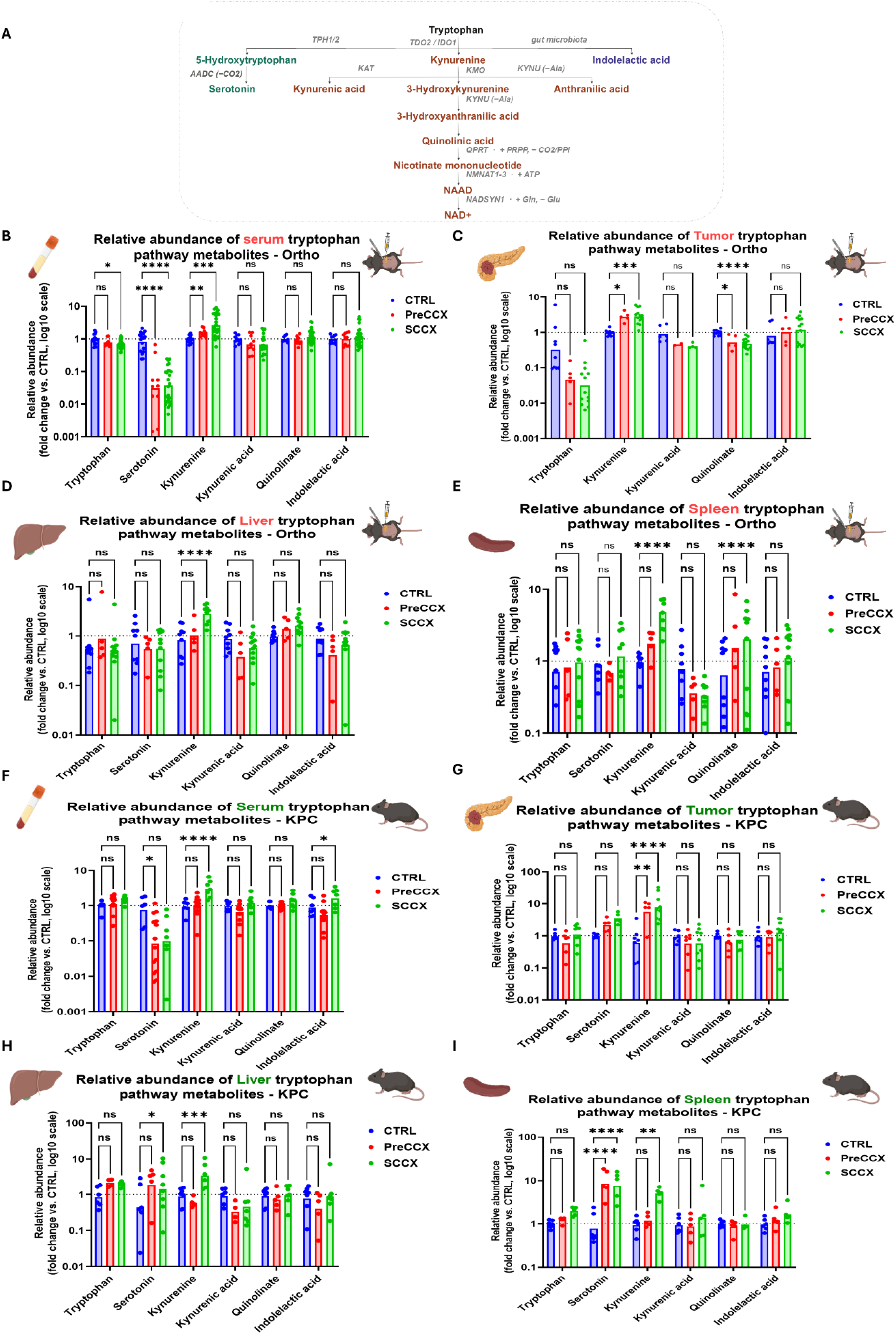
Tryptophan-pathway metabolites across PDAC cachexia models. (A) Tryptophan pathway. (B–E) Metabolites in serum (B), pancreas/tumor (C), liver (D), and spleen (E) of CTRL, PreCCX, and SCCX (Figure 1A) orthotopic PDAC mice; fold change versus CTRL (log10 scale). *n* = 18/10/26 (B), 9/5/13 (C), 9/5/12 (D), and 9/5/13 (E) CTRL/PreCCX/SCCX mice. (F–I) Same for GEMM KPC mice: serum (F), pancreas/tumor (G), liver (H), and spleen (I). *n* = 7/14/8 (F), 7/7/9 (G), 7/7/8 (H), and 6/5/5 (I) mice. *n* varies by metabolite. Data are mean ± SEM; two-way repeated-measures ANOVA/mixed-effects model with Geisser–Greenhouse correction and Dunnett’s test versus CTRL. \**p* < 0.05, \*\**p* < 0.01, \*\*\**p* < 0.001, \*\*\*\**p* < 0.0001. See also Figures S3–S6.

Tumors showed a partially conserved metabolic fingerprint dominated by creatine and creatine phosphate depletion, consistent with energetic strain. Model-specific changes included increased fumarate and 2-hydroxybutyric acid in orthotopic severe cachexia tumors and decreased α-ketoglutarate and isocitrate in KPC tumors. PCA reveals separated tumor from control pancreas in both models, with partial overlap (Figures S3C and S3D; Figures S5C and S5D). The liver showed the most extensive remodeling with increased 3-hydroxybutyrate in the orthotopic model, consistent with enhanced hepatic ketogenesis. This rewiring also extended to amino acid metabolism, exemplified by altered phenylalanine in cachectic livers, consistent with broader perturbation of amino acid handling in cachexia^16,42,43^. Metabolites linked to creatine-phosphocreatine buffering were also altered, suggesting disruption of cellular energy-buffering pathways under heightened energetic demand^44,45^. PCA showed progressive separation of cachectic from control livers in both models, with pre-cachectic samples intermediate (Figures S4A and S4B; Figures S6A and S6B).

In orthotopic mice, the spleen showed substantial remodeling, including increased phenylalanine, cystine, glutamine and creatine, while creatine phosphate decreased, consistent with altered creatine–phosphocreatine energy buffering (Figures S4C and S4D). KPC spleen similarly showed increased isoleucine, glutamine, and valine and a distinct PCA profile (Figures S6C and S6D). Notably, splenic serotonin followed the opposite trajectory of every other tissue examined, rising rather than falling in the KPC model (Figure 3I). Given the pronounced splenomegaly in KPC mice, and that platelets are the dominant peripheral serotonin reservoir and are sequestered in the spleen, one possible explanation is platelet pooling secondary to splenomegaly^46,47^, rather than a genuine reversal of splenic tryptophan flux, and further studies are needed to confirm this.

Together, these findings support a coordinated, tumor orchestrated metabolic program in pancreatic cancer cachexia, marked by systemic tryptophan diversion toward kynurenine and impaired energy buffering across tissues.

Our findings are supported by human cancer data, though these examined individual pathway components of tryptophan metabolism in separate disease settings. In human PDAC, kynurenine is elevated in tumors versus patient-matched adjacent pancreas, ranking among the most increased of 127 metabolites in 49 patients^48^. Kynurenine is likewise raised in renal and colon tumors versus normal tissue^49,50^, and circulating kynurenine is elevated in lung, gynecologic, and hepatocellular cancers, tracking with stage and predicting survival^51–53^. In paired human rectal tumors, kynurenine rose while serotonin and 5-HIAA fell, showing both branches shift within the same tumor^54^. Fragmentary evidence links the pathway to cachexia: serum tryptophan is depleted in cachectic patients and correlates inversely with immune activation^55^, and kynurenine pathway activation has separately been associated with fatigue and depressed mood in solid tumors^56^. Each captures a single tissue or symptom. Our data supply that link, showing tryptophan is coordinately redirected toward kynurenine and away from serotonin across multiple organs during cachexia, and that this shift accompanies both wasting and behavioral features of the syndrome.

### Transcriptional evidence of systemic kynurenine pathway activation in cachexia

The metabolic bottleneck observed across serum, tumor, liver, and spleen, with kynurenine accumulation but no increase in downstream metabolites, raised the question, does this phenotype reflect increased kynurenine production, impaired downstream catabolism, or both? Transcriptomic profiling across tumor, liver, adipose, and muscle showed that kynurenine synthesis genes *Ido1* and *Afmid* were markedly upregulated in tumors, whereas downstream catabolic enzymes *Kmo, Kynu, Haao,* and *Qprt* were unchanged or reduced, and serotonin pathway genes (*Tph1, Ddc, Maoa,* and *Slc6a4*) were differentially regulated across tissues and models (Figure 4A).

**Figure 4.**
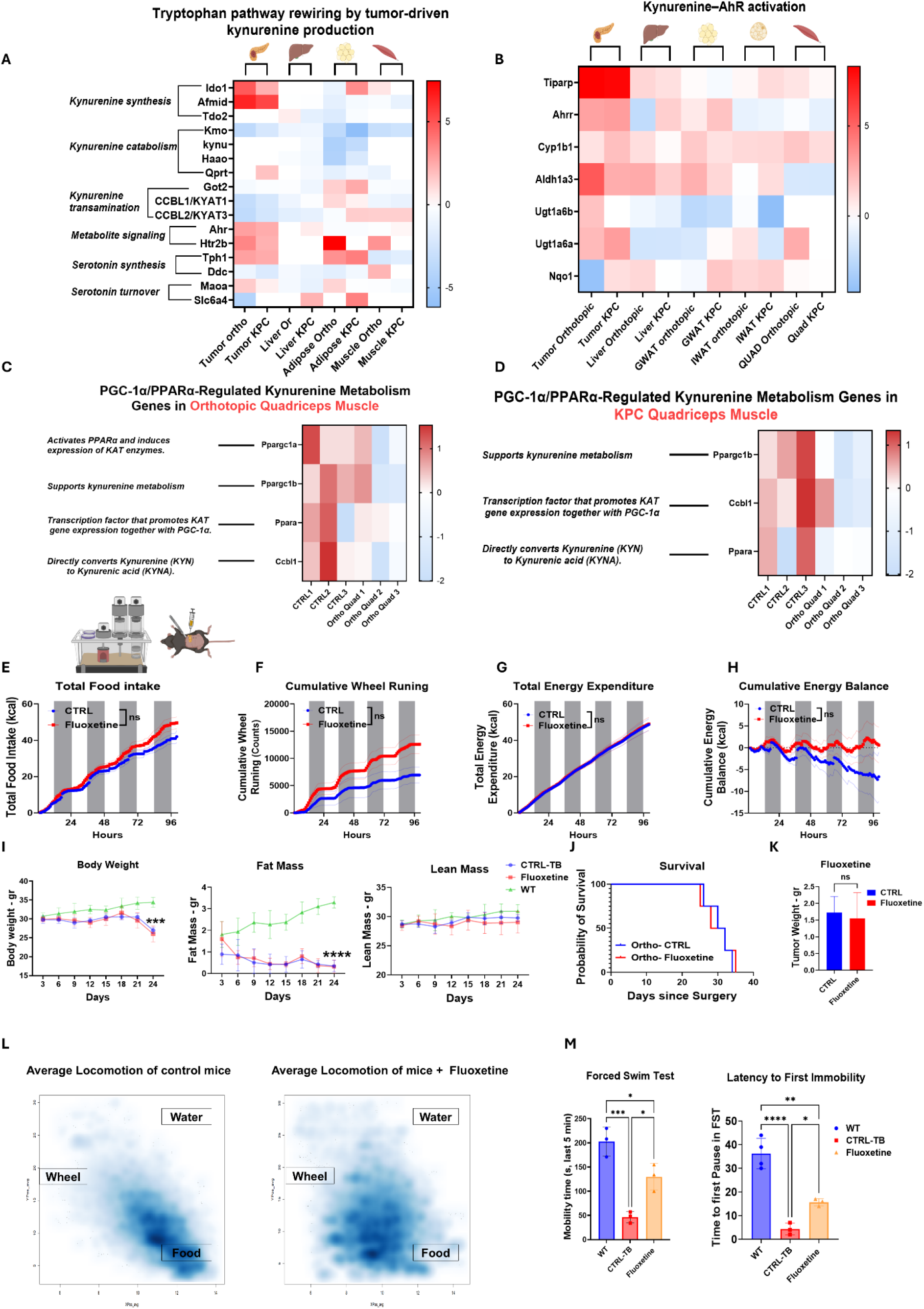
Tryptophan metabolism and fluoxetine responses in PDAC cachexia. (A and B) Tryptophan-pathway genes (A) and aryl hydrocarbon receptor (AhR) targets (B) across tissues of orthotopic PDAC and GEMM KPC mice. GWAT and IWAT, gonadal and inguinal white adipose tissue. (C and D) PGC-1α/PPARα-regulated kynurenine-metabolism genes in quadriceps of orthotopic PDAC (C) and GEMM KPC (D) mice. *n* = 3 mice per group (A–D). (E–H) Metabolic-cage analysis of mixed-sex orthotopic PDAC-bearing mice given vehicle (CTRL) or fluoxetine: food intake (E), wheel running (F), energy expenditure (G), and energy balance (H). *n* = 4 (CTRL) and 6 (fluoxetine) mice. (I) Body weight, fat mass, and lean mass in WT and tumor-bearing mice treated with vehicle (CTRL-TB) or fluoxetine. *n* = 5 per group. (J–L) Survival (J), tumor weight (K), and cage-occupancy maps (L) of CTRL and fluoxetine-treated mice. *n* = 4 (J), 5 (K), and 3 (L) per group. (M) Forced swim test in WT, CTRL-TB, and fluoxetine-treated mice. *n* = 4, 3, and 3 mice, respectively. Data are mean ± SEM; two-way repeated-measures ANOVA/mixed-effects model with Geisser–Greenhouse correction and Šídák’s test (E–H), two-way repeated-measures ANOVA with multiple comparisons (I), log-rank test (J), unpaired Student’s *t* test (K), or one-way ANOVA with Tukey’s test (M). \**p* < 0.05, \*\**p* < 0.01, \*\*\**p* < 0.001, \*\*\*\**p* < 0.0001. See also Figure S7.

Indoleamine 2,3-dioxygenase 1 (IDO1) was significantly elevated in both KPC and orthotopic tumors versus normal pancreas, with stronger induction in the orthotopic model (Figure S7A), accompanied by elevated *Il6* in orthotopic tumors (Figure S7B) and robust hepatic *Il6* induction in cachectic KPC livers relative to controls (Figure S7C), consistent with a model in which IL-6, a known inducer of IDO1^57^, sustains kynurenine pathway activation both locally and systemically.

Kynurenine also functions as an endogenous ligand for the aryl hydrocarbon receptor (AhR), a transcription factor coupling metabolic cues to immune and stress-response gene programs^58,59^. We therefore asked whether kynurenine accumulation was transcriptionally active rather than metabolically inert. Canonical AhR targets (*Tiparp*, *Ahrr*, *Cyp1b1*, *Aldh1a3*, *Ugt1a6a*, *Ugt1a6b*) were coordinately induced across tumor, liver, gonadal and inguinal white adipose tissue (GWAT, IWAT), and quadriceps muscle (Quad) in both models (Figure 4B), consistent with kynurenine ligand-driven AhR activation^60,61^. Induction was strongest in tumor but clear in liver and both adipose depots, and included the AhR-dependent detoxification genes *Ugt1a6a/b*. Thus, these data support a model in which tumor-derived inflammation, particularly IL-6-driven IDO1 induction, drives a systemic program favoring kynurenine production and AhR signaling over serotonin synthesis.

Unchanged kynurenic acid, quinolinate, and indolelactic acid pointed to impaired metabolism downstream of production. Agudelo et al. described a skeletal muscle PGC-1α/PPARα–KAT program converting kynurenine to kynurenic acid^62^. Orthotopic quadriceps showed reduced expression of *Ppargc1a*, *Ppargc1b*, *Ppara*, and *Ccbl1* (*Kyat1*) (Figure 4C), and KPC quadriceps a similarly coordinated reduction in *Ppargc1b*, *Ppara*, and *Ccbl1* (Figure 4D), indicating impaired muscle conversion of kynurenine to kynurenic acid, a tissue intrinsic mechanism compounding the systemic bottleneck.

### Conserved alteration of tryptophan metabolism across distinct cancer cachexia models

To test whether this rewiring extends to other tumor types, we examined two genetically engineered lung cancer cachexia models, KP (Kras^LSL-G12D/+^; p53^fl/fl^) and KL (Kras^LSL-G12D/+^; Lkb1^fl/fl^) (Figures S7D and S7E). Consistent with our pancreatic models, both lung cancer models showed reduced serotonin with increased kynurenine, though affected tissues differed. KP mice showed increased serum and liver kynurenine with reduced serum serotonin, and KL mice showed reduced serum serotonin with increased liver kynurenine. Kynurenic acid was reduced or unchanged despite rising kynurenine in both models, mirroring the bottleneck identified in PDAC. These findings support kynurenine accumulation with reduced serotonin as a conserved feature of cancer cachexia across genetically distinct tumor models, suggesting this tryptophan pathway alteration may be a common systemic response extending beyond PDAC.

### Cellular energy crisis: another conserved feature of pancreatic cancer cachexia

The altered creatine–phosphocreatine pattern observed by metabolomics suggested broader disruption of cellular energy homeostasis, prompting us to examine whether cellular energy status was perturbed across tissues. We therefore calculated adenylate energy charge, an index of the phosphorylation state of the adenine nucleotide pool, in soleus muscle, liver, and spleen across disease stages. Energy charge declined progressively from control to pre-cachectic to severely cachectic animals in all three tissues (Figure S7F), demonstrating a multi-organ reduction in cellular energy status. Because energy charge can be influenced by changes in energy supply, energy demand, or both, these data do not distinguish which component primarily drives the observed decline. These findings identify impaired cellular energy homeostasis as a conserved feature of pancreatic cancer cachexia.

### Restoration of serotonin signaling improves activity and appetite but does not prevent wasting and lethality

Reduced serotonin signaling has been strongly associated with depressive states, and depression is frequently observed in patients with pancreatic cancer cachexia^63,64^. Moreover, increased kynurenine production via the tryptophan degrading enzyme IDO1 has been reported to induce depression-like behavior in mice and humans^65,66^. Together, this suggests tumor-driven alteration of tryptophan metabolism may contribute to neurobehavioral features of cachexia.

To test whether restoring serotonin signaling could improve these features, we treated orthotopic tumor bearing mice with fluoxetine, a selective serotonin reuptake inhibitor (SSRI) that raises synaptic serotonin levels by blocking its reuptake transporter, SERT^67^. Fluoxetine increased cumulative food intake and, most prominently, voluntary wheel running (Figures 4E and 4F), consistent with restoration of motivated behavior rather than generalized movement. Video monitoring showed fluoxetine-treated mice moving freely through the cage compared with the hunched, immobile posture of controls, consistent with reduced depression-like behavior (Video S1). Total energy expenditure remained unchanged (Figure 4G). Cumulative energy balance remained near neutral with fluoxetine but progressively declined in controls (Figure 4H), consistent with increased food intake offsetting ongoing energetic demands.

Despite this, fluoxetine-treated mice showed no preservation of body weight, fat mass, or lean mass (Figure 4I), and survival (Figure 4J) and tumor burden (Figure 4K) were unchanged. Thus, improved feeding, voluntary activity, and energy balance were insufficient to prevent wasting or extend survival, separating reversible behavioral and energetic defects from the lethal wasting phenotype.

We next tested this response in KPC mice implanted after cachexia onset with sustained release fluoxetine pellets (∼0.37 mg/day over 30 days, comparable to the orthotopic dose). Fluoxetine similarly increased cumulative food intake and voluntary wheel running, while total energy expenditure remained unchanged, resulting in a more positive cumulative energy balance (Figures S7G–S7J) and recapitulating the energetic benefit observed in the orthotopic model. Again, fluoxetine did not preserve body weight, fat mass, or lean mass or improve survival or tumor burden (Figures S7K–S7M), reproducing the dissociation between improved feeding, activity and persistent wasting.

Serum metabolomics showed increased circulating serotonin, supporting engagement of the serotonergic axis (Figure S7U). Together, these findings demonstrate a reproducible response across two PDAC cachexia models: fluoxetine increased food intake and voluntary activity and improved whole body energy balance but did not prevent wasting or prolong survival, showing that neurobehavioral dysfunction and negative energy balance are pharmacologically reversible and uncoupled from the lethal wasting aspect of cachexia.

### Bypassing serotonin biosynthesis with 5-HTP has limited effects on PDAC cachexia

To test whether enhancing serotonin synthesis could alleviate cachexia-related symptoms, we supplemented the diet with 5-hydroxytryptophan (5-HTP), the immediate serotonin precursor that readily crosses the blood-brain barrier^68,69^, thereby bypassing upstream tryptophan metabolism and kynurenine diversion. 5-HTP did not alter food intake but modestly increased voluntary wheel running (Figures S7N and S7O). Total energy expenditure was modestly increased, whereas cumulative energy balance remained negative (Figures S7P and S7Q). 5-HTP did not preserve body weight, fat mass, or lean mass, or alter tumor burden or survival (Figures S7R–S7T). Thus, 5-HTP produced only a modest increase in activity without improving energy balance or wasting.

### Serotonergic modulation of depression-like behavior in pancreatic cancer cachexia

Pancreatic cancer cachexia is accompanied by behavioral changes consistent with depression-like states, including reduced activity and loss of motivation^38,70^. Because fluoxetine increased appetite and voluntary activity, we hypothesized that altered serotonin signaling contributes to these deficits. Spatial analysis of movement in metabolic cages showed that control cachectic mice remained near the food source, whereas fluoxetine-treated mice moved throughout the cage, including around food, water, and wheel areas (Figure 4L), consistent with increased exploration and reduced behavioral restriction.

To further assess depression-like behavior, we performed the forced swim test, in which increased immobility is widely used as an index of depression-like behavior^71^. Control non-tumor bearing mice readily swam, tumor bearing mice receiving placebo pellets rapidly became immobile, spending most of the test floating rather than actively swimming, consistent with increased depression-like behavior (Figure 4M). In contrast, fluoxetine-treated mice swam significantly longer and showed delayed immobility, consistent with reduced behavioral despair and improved coping behavior (Video S2). Although they remained less mobile than non-tumor bearing controls, their improvement over placebo-treated tumor bearing mice indicates partial rescue of behavioral function, supporting that cachexia-associated depression-like behavior is alleviated by enhanced serotonergic signaling.

### Targeting the kynurenine pathway increases feeding and locomotor activity without altering tissue wasting and lethality

To test whether targeting the kynurenine branch could alleviate cachexia symptoms, we inhibited IDO1 with the selective inhibitor Epacadostat^72,73^. IDO1 is a rate-limiting enzyme that catalyzes tryptophan degradation to kynurenine^74^. Cachectic mice received Epacadostat (100 mg/kg by oral gavage) and underwent Promethion monitoring and longitudinal EchoMRI. IDO1 inhibition markedly increased food intake (Figure 5A) and increased voluntary wheel running without reaching significance (Figure 5B). Energy expenditure was modestly elevated but not significant (Figure 5C), whereas cumulative energy balance improved significantly (Figure 5D). Body weight, fat mass, lean mass, survival, and tumor burden were unchanged (Figures 5E–5G). Serum metabolomics confirmed reduced circulating kynurenine in Epacadostat-treated mice (Figure S7V), consistent with pathway inhibition. Thus, IDO1 inhibition improved feeding and energy balance, with trends toward greater activity, without altering tissue wasting or disease outcome, further dissociating the behavioral, energetic, and lethal arms of PDAC cachexia.

**Figure 5.**
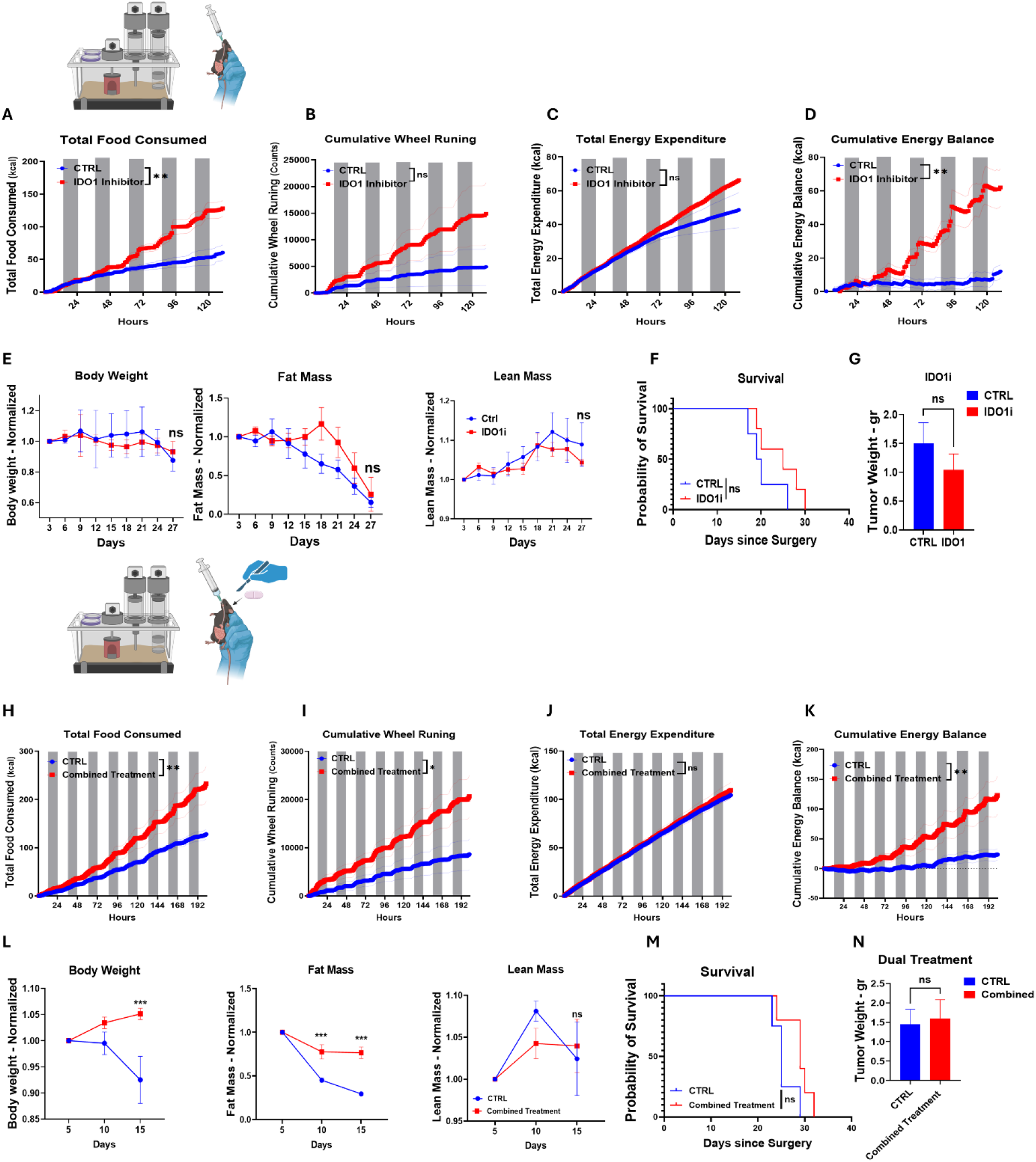
IDO1 inhibition and dual treatment in PDAC cachexia. (A–D) Metabolic-cage analysis of male orthotopic PDAC-bearing mice given vehicle (CTRL) or the IDO1 inhibitor epacadostat (IDO1i): food intake (A), wheel running (B), energy expenditure (C), and energy balance (D). *n* = 4 (CTRL) and 3 (IDO1i) mice. (E–G) Body weight, fat mass, and lean mass normalized to baseline (E), survival (F), and tumor weight (G) of vehicle- and IDO1i-treated mice. *n* = 5 per group. (H–K) Metabolic-cage analysis of male orthotopic PDAC-bearing mice given vehicle (CTRL) or fluoxetine plus IDO1i (dual treatment): food intake (H), wheel running (I), energy expenditure (J), and energy balance (K). *n* = 7 (CTRL) and 5 (dual) mice. (L–N) Body weight, fat mass, and lean mass normalized to baseline (L), survival (M), and tumor weight (N) of vehicle- and dual-treated mice. *n* = 4 per group. Data are mean ± SEM; two-way repeated-measures ANOVA/mixed-effects model with Geisser–Greenhouse correction and Šídák’s test (A–E and H–L), log-rank test (F and M), or unpaired Student’s *t* test (G and N). \**p* < 0.05, \*\**p* < 0.01, \*\*\**p* < 0.001, \*\*\*\**p* < 0.0001. See also Figure S7.

### Combined targeting of tryptophan metabolism improves activity and preserves fat mass in cachexia

To determine whether combined serotonergic and IDO1 targeting could more effectively alleviate cachexia, we treated cachectic mice with sustained release fluoxetine pellet and IDO1 inhibitor. Combined treatment significantly increased food intake and voluntary wheel running (Figures 5H and 5I), indicating improved feeding and spontaneous activity. Total energy expenditure was unchanged (Figure 5J), whereas cumulative energy balance was significantly more positive (Figure 5K). Unlike either single agent, combined treatment preserved body composition: treated mice maintained significantly greater body weight and fat mass, whereas lean mass remained unchanged (Figure 5L). However, survival and tumor burden (Figures 5M and 5N) remained unchanged. Thus, simultaneous targeting of both branches improved feeding, activity, and energy balance and, translated these gains into preservation of body weight and fat mass without altering tumor burden or survival.

### Stable isotope tracing reveals kynurenine pathway activation as the dominant fate of tryptophan in PDAC

Because steady-state metabolomics cannot resolve pathway utilization, we performed in vivo U-¹³C₁₁-tryptophan tracing to track newly available tryptophan through the serotonin, kynurenine, and indole pathways (Figure 6A). Serum tracer enrichment was modestly higher in PDAC-bearing mice (Figure 6B). To correct for differences in systemic tracer exposure, tissue metabolite labeling was normalized to matched serum enrichment from the same animal. Serum normalized labeled tryptophan was unchanged in pancreatic/tumor and liver (Figure 6C), indicating similar precursor labeling relative to systemic exposure. In contrast, labeled kynurenine was markedly increased in tumor and liver (Figure 6D), demonstrating preferential channeling of newly available tryptophan into the kynurenine pathway. This did not extend uniformly downstream: labeled kynurenic acid remained unchanged in both tissues (Figure 6E). Labeled 5-HTP was unchanged in pancreatic/tumor tissue but increased in liver (Figure 6F), while labeled serotonin remained low and unchanged in both (Figure 6G), revealing a reciprocal suppression of the serotonin branch at its immediate precursor. Labeled indolelactic acid was unchanged in tumor but increased in liver (Figure 6H), indicating branch and tissue selective tracer redistribution rather than globally increased tracer availability. Thus, enhanced tryptophan to kynurenine conversion is the dominant remodeling of tryptophan metabolism in PDAC, with reduced serotonin pathway engagement and limited downstream conversion of the expanded kynurenine pool.

**Figure 6.**
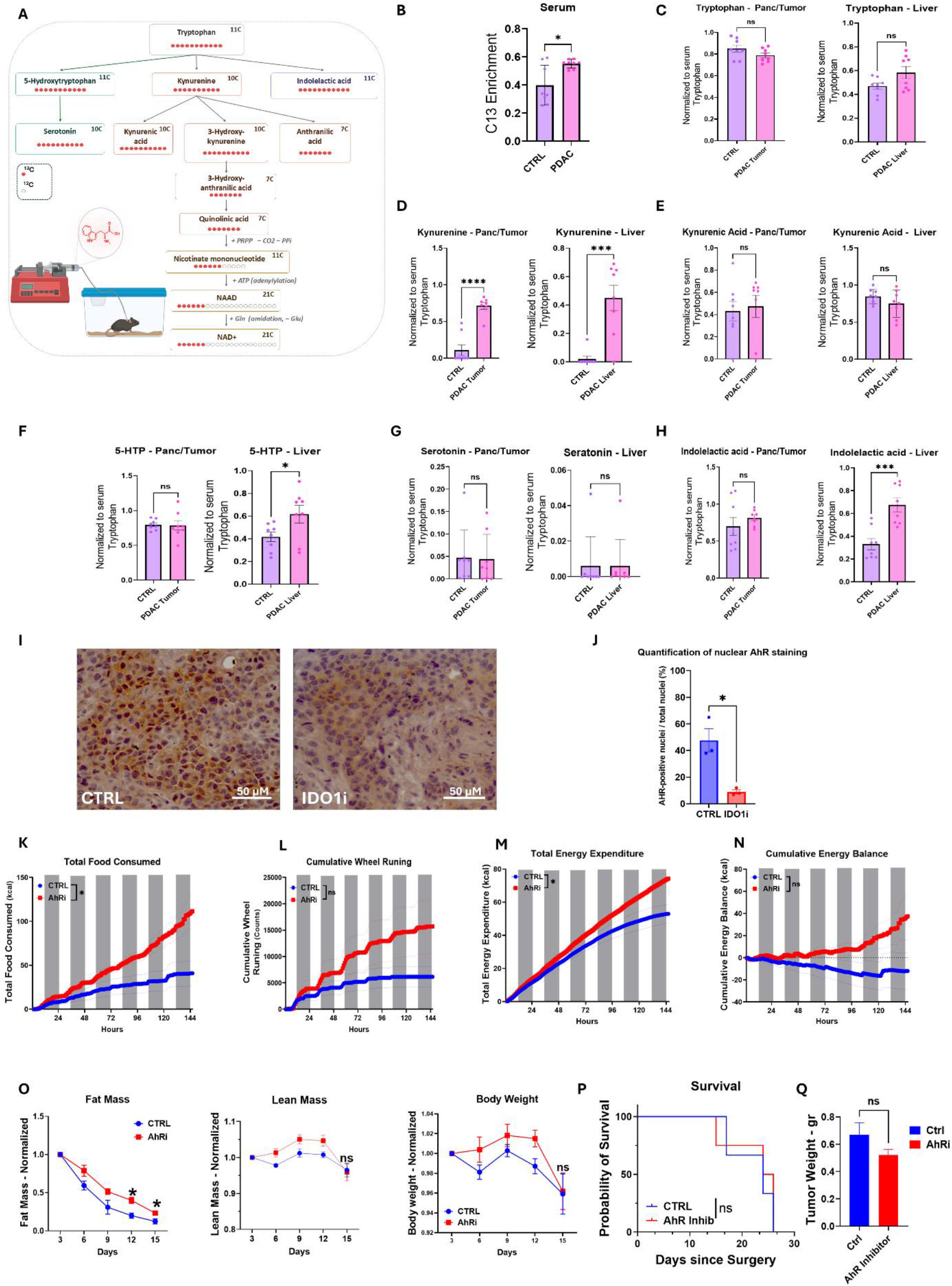
Tryptophan tracing and AhR modulation in PDAC cachexia. (A) In vivo U-^13^C_11_-tryptophan tracing. (B–H) Serum ^13^C enrichment (B) and labeling of tryptophan (C), kynurenine (D), kynurenic acid (E), 5-hydroxytryptophan (F), serotonin (G), and indolelactic acid (H) in pancreas/tumor and liver (normalized to serum tryptophan) of control and orthotopic PDAC mice. *n* = 8 per group. (I and J) Representative aryl hydrocarbon receptor (AhR) immunohistochemistry (I) and percentage of AhR-positive nuclei (J) in orthotopic tumors from vehicle- and IDO1 inhibitor (IDO1i)-treated mice. Scale bars, 50 μm. *n* = 3 per group. (K–N) Metabolic-cage analysis of male orthotopic PDAC-bearing mice given vehicle (CTRL) or the AhR antagonist CH-223191 (AhRi): food intake (K), wheel running (L), energy expenditure (M), and energy balance (N). *n* = 5 (CTRL) and 6 (AhRi) mice. (O–Q) Fat mass, lean mass, and body weight normalized to baseline (O), survival (P), and tumor weight (Q) of vehicle- and AhRi-treated mice. *n* = 4 per group. Data are mean ± SEM; unpaired Student’s *t* test (B–H, J, and Q), two-way repeated-measures ANOVA/mixed-effects model with Šídák’s test (K–O; Geisser–Greenhouse correction for K–N), or log-rank test (P). \**p* < 0.05, \*\**p* < 0.01, \*\*\**p* < 0.001, \*\*\*\**p* < 0.0001. See also Figure S7.

### AhR inhibition functionally phenocopies disruption of the tryptophan metabolism pathway in cachectic PDAC

PDAC cachexia showed molecular features consistent with AhR pathway engagement, including tissue-specific alterations in canonical AhR-responsive genes (Figure 4B). AhR is a ligand-dependent transcription factor activated by endogenous metabolites, including kynurenine, with activation promoting nuclear translocation and transcriptional activity^75^. Because tracing identified kynurenine as a predominant metabolic fate of tryptophan in PDAC cachexia, we asked whether this metabolic rewiring was associated with downstream AhR activation in tumors. Because IDO1 converts tryptophan to kynurenine, we examined tumor AhR localization after IDO1 inhibition using an anti-AhR antibody previously used for immunohistochemistry^76,77^. Vehicle-treated tumors showed spatially heterogeneous AhR staining, with discrete tumor cell clusters displaying nuclear localization. IDO1 inhibition markedly reduced these nuclear AhR positive foci (Figure 6I), as confirmed by quantitative analysis (Figure 6J). This links IDO1 dependent kynurenine production to AhR activation in tumors, while allowing for contributions from other endogenous AhR ligands. To test whether AhR signaling functionally contributes to cachexia, orthotopic PDAC-bearing mice received the AhR antagonist CH-223191 by oral gavage. AhR inhibition significantly increased food intake (Figure 6K) and raised voluntary wheel running, although not significantly (Figures 6L). Total energy expenditure increased significantly (Figure 6M), while cumulative energy balance shifted positively (Figure 6N). AhR inhibition also significantly attenuated fat mass loss, whereas lean mass and body weight remained unchanged (Figure 6O). Survival and tumor weight were unaffected (Figures 6P and 6Q). Thus, AhR blockade reproduced key effects of upstream kynurenine pathway inhibition, improving feeding, energy balance, and adiposity without altering tumor burden or survival. These findings position AhR as a functional downstream mediator of host metabolic and behavioral responses to altered tryptophan metabolism rather than a determinant of tumor growth or lethality.

### Excess dietary tryptophan promotes kynurenine production only in PDAC and enhances lethal cachexia

To determine whether increased substrate availability could overcome altered tryptophan metabolism, non-tumor bearing and orthotopic PDAC-bearing mice were randomized on day 5 to control or 10× tryptophan diets and analyzed on day 12 and at the cachectic endpoint (Figure 7A). Supplementation significantly shortened survival in PDAC-bearing mice (Figure 7B), indicating that added tryptophan was detrimental rather than corrective. We therefore performed serum metabolomic profiling to define its effects on circulating tryptophan metabolites (Figure 7C).

**Figure 7.**
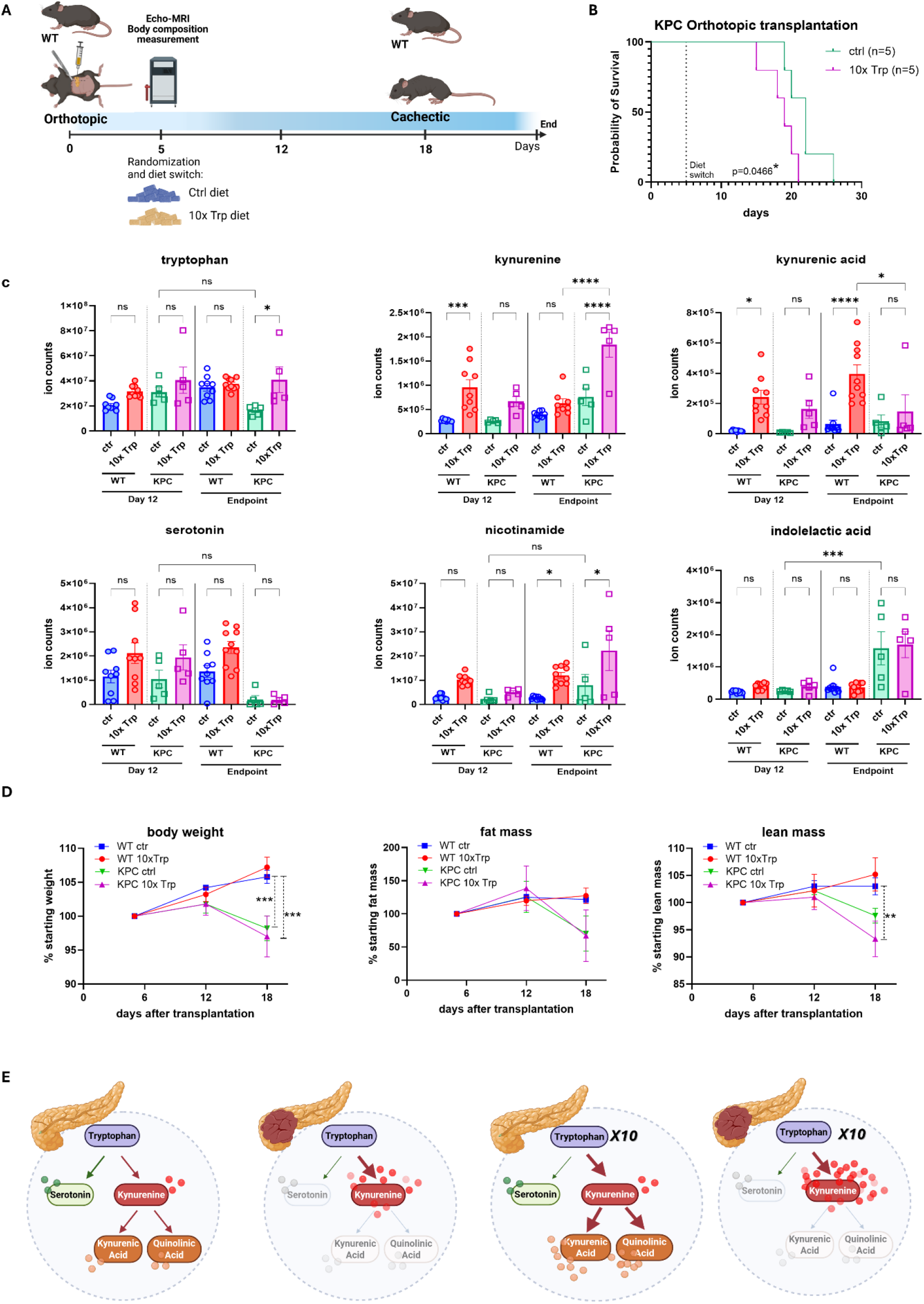
Dietary tryptophan enrichment in PDAC cachexia. (A) WT and KPC-orthotopic mice switched on day 5 to control or 10× tryptophan (Trp) diet. (B) Survival of KPC-orthotopic mice. *n* = 5 per group. (C) Serum tryptophan-pathway metabolites at day 12 and endpoint. *n* = 9/9 (day 12) and 9/10 (endpoint) control/10× Trp WT and 5 KPC mice per group. (D) Body weight, fat mass, and lean mass relative to baseline. *n* = 5 mice per group. (E) Summary of tryptophan-pathway changes. Data are mean ± SEM; log-rank test (B) or ordinary one-way (C) or two-way (D) ANOVA with Tukey’s test. \**p* < 0.05, \*\**p* < 0.01, \*\*\**p* < 0.001, \*\*\*\**p* < 0.0001.

Despite a 10-fold dietary increase, circulating tryptophan rose only modestly, reaching significance only in PDAC-bearing mice at endpoint. At day 12, supplementation increased kynurenine and kynurenic acid in non-tumor bearing mice, whereas neither changed significantly in PDAC-bearing mice. By endpoint, non-tumor bearing mice accumulated kynurenic acid without sustained kynurenine elevation, whereas PDAC-bearing mice showed marked kynurenine accumulation exceeding both diet matched PDAC controls and supplemented non-tumor bearing mice, without a rise in kynurenic acid. Circulating serotonin remained unchanged, indicating that added tryptophan did not restore the serotonergic pool. Nicotinamide increased at endpoint in both groups, whereas indolelactic acid was unchanged.

Tryptophan supplementation also failed to preserve body weight, fat mass, or lean mass in PDAC-bearing mice (Figure 7D). This complements the tracing data, which showed preferential incorporation of newly available tryptophan into kynurenine with limited downstream labeling. Together, they show that excess tryptophan does not normalize metabolism in PDAC cachexia but instead reveals differential partitioning toward kynurenic acid in healthy mice and kynurenine in tumor bearing mice (Figure 7E).

## Discussion

Pancreatic cancer cachexia extends beyond anorexia and tissue wasting. Across PDAC mouse models, we identify three largely separable axes: tumor to periphery inflammatory signaling converging on the liver, a conserved multi-organ decline in cellular energy charge, and systemic tryptophan remodeling from serotonin toward the kynurenine–AhR axis. Each was independently targetable, yet every intervention corrected only a subset of the phenotype, reframing PDAC cachexia as coordinated, partially overlapping catabolic programs downstream of the tumor.

Inflammatory and tumor-derived signals induce IDO1 and tryptophan 2,3-dioxygenase (TDO)^78,79^, diverting tryptophan from serotonin toward kynurenine^80,81^; its contribution to the systemic manifestations of cancer, as opposed to antitumor immunity, has remained less clear^58,79^. In this context, our data indicate an active, tumor-conditioned program rather than a passive consequence of malignancy: tumors selectively upregulated *Ido1* and *Afmid* without inducing downstream catabolic genes/enzymes, while isotope tracing showed tryptophan channeled preferentially into kynurenine rather than serotonin, with limited conversion to kynurenic acid, quinolinate, or indolelactic acid. This bottleneck recurred across tumor, liver, and adipose tissue in two PDAC and two lung cancer models (KP and KL), supporting a conserved metabolic feature of cancer cachexia.

Serotonin regulates feeding, mood, and locomotion, all impaired in advanced cancer and cachexia. Inflammation and altered serotonergic signaling are linked to anorexia and neurobehavioral dysfunction in chronic disease and cancer^82,83^, and increased tryptophan catabolism to depression and fatigue in patients with cancer^56,84,85^. Accordingly, serotonin was reduced in cachectic mice, and serotonergic restoration improved food intake, activity, and depression-like behavior. Across interventions, however, improved intake and energy balance remained dissociated from wasting: fluoxetine, 5-HTP, IDO1 inhibition, and AhR antagonism each improved feeding, activity, and/or energy balance, yet none slowed body mass loss or extended survival. PDAC cachexia is therefore driven not by insufficient caloric intake but by catabolic programs that persist despite improved energy balance. Only AhR antagonism and combined fluoxetine plus IDO1 inhibition preserved fat mass, indicating partial but incomplete control of wasting.

Tracing further showed increased kynurenine flux without proportional increases downstream, most evident with dietary tryptophan. In tumor free mice, excess tryptophan increased kynurenic acid instead of kynurenine, consistent with further metabolism of kynurenine into a metabolite with limited blood–brain barrier permeability^86,87^. In PDAC-bearing mice, supplementation instead caused marked kynurenine accumulation without increased kynurenic acid. The tumor bearing state thus shifts kynurenine handling toward accumulation rather than kynurenic acid production, relevant given the differential blood–brain barrier permeability of these metabolites^86^. The high tryptophan diet also accelerated mortality in PDAC-bearing mice, showing that added substrate exacerbates rather than corrects kynurenine pathway dysregulation.

Increased kynurenine pathway activity has been associated with fatigue, reduced physical performance, muscle dysfunction, and neurobehavioral abnormalities in inflammatory and cancer-associated conditions^56,88^, and IDO1 has been pursued as an anticancer target because tryptophan depletion and kynurenine production suppress antitumor immunity^78,89^. Although IDO1 inhibition alone primarily improved feeding and activity, combining it with serotonin restoration also preserved adipose mass, suggesting that targeting of both tryptophan branches may better address the behavioral and metabolic features of cachexia.

One downstream mechanism linking kynurenine to cachexia is AhR activation. Kynurenine acts as an endogenous AhR ligand, connecting tryptophan metabolism to transcriptional programs of immune regulation, inflammation, and cellular metabolism^78,90,91^. IDO1 inhibition altered AhR subcellular localization, consistent with reduced pathway activation when kynurenine production was suppressed, in line with evidence that elevated AhR activity driven by kynurenine pathway metabolites promotes cancer progression and wasting, as in gastric adenocarcinoma, where AhR overexpression drove tumor growth and its suppression alleviated cachexia^92^. Consistent with our findings, Leng and colleagues showed that dietary tryptophan promotes multiple myeloma engraftment through IDO1-dependent catabolism and host myeloid AhR signaling. In our model, Direct AhR inhibition improved food intake, locomotion, and fat mass preservation, establishing AhR as a functional mediator of selected cachexia phenotypes rather than merely a marker of altered kynurenine metabolism. It did not reverse all manifestations of cachexia or improve survival, indicating that AhR drives a defined subset of host abnormalities rather than the whole syndrome.

Together, these findings establish altered tryptophan partitioning, with reduced serotonergic output and kynurenine–AhR activation, as an active, targetable component of PDAC cachexia rather than a passive biomarker. They further suggest that depression, anorexia, and inactivity in cancer cachexia have a biochemical basis that may be therapeutically addressed with available agents to improve quality of life. With the accompanying study, our findings highlight tryptophan metabolism as a therapeutic avenue for improving key manifestations of cancer cachexia.

### Limitations of the study

Our behavioral assays interrogate selected aspects of locomotion, motivation and depression-like behavior but not the full neurobehavioral spectrum of cachexia. We did not measure central tryptophan, kynurenine, or serotonin, so their contribution beyond peripheral signaling remains inferred. Our conclusions rest on pharmacologic rather than genetic manipulation of IDO1, AhR and serotonin signaling. Without tissue specific loss or gain of function models, off-target drug effects cannot be fully excluded. Finally, the impaired hepatic AhR signaling reported previously differs from the tumor associated AhR activation seen here, a discrepancy that may reflect tissue context, tumor model, or disease stage and warrants direct comparative investigation.

## Resource availability

Raw data supporting the main and supplementary figures are provided as source data files. Additional datasets, analyses, and methodological details are available from the lead contact, EW, upon request.

## Acknowledgments

This work was supported by Cancer Grand Challenges CGCATF-2021/100026 (CRUK) and NCI grant 1OT2CA278609-01 (NCI) to E.W., M.D.G., and T.J., the Ludwig Institute for Cancer Research Ludwig Princeton Branch and NIH grant R01CA163591 to E.W. and J.D.R. J.Y.G. was supported by NIH grants R01CA237347 and R01CA308211. E.P. was supported by postdoctoral fellowship COCR26PDF010 from the New Jersey Commission on Cancer Research (NJCCR). M.G.J. was also supported by 1T32CA257957. T.G.A. was supported by NIH grant DK109714. We acknowledge support from the Rutgers Cancer Institute Metabolomics Shared Resource (NCI-CCSG P30CA072720-6852) and the Biospecimen Repository and Histopathology Service Shared Resource (P30CA072720-5919). We thank the Dr. Gulhati laboratory at Rutgers Cancer Institute for providing the PDAC cell line, Dr. Mona Arabzadeh for bioinformatic support, and Drs. Shin-Heng Chiou and Sanith Cheriyamundath for guidance on PDAC models. We also thank members of the White laboratory for their support. Schematics were created with BioRender.com.

## Author Contributions

D.F. designed and performed experiments, analyzed and interpreted the data, and wrote the manuscript. E.P. performed the dietary tryptophan supplementation experiments. M.G.J. contributed to isotope-tracing analyses and to the analysis of data from the KP and KL mouse models. M.I. and T.D. contributed to metabolic phenotyping experiments and data analysis. C.B. assisted with forced swim test experiments. S.L.D. and B.J.R. contributed to single-nucleus RNA-sequencing analysis. J.P. and Z.H. assisted with animal husbandry and genotyping, and Z.H. performed jugular vein catheterization surgeries. L.S., C.L., and C.G. assisted with mouse experiments, daily feeding, tissue collection, and data acquisition. T.G.A., J.D.R., J.Y.G., M.D.G., and T.J. provided scientific input and guidance. E.W. conceived and supervised the study. All authors reviewed, edited, and approved the manuscript.

## STAR METHODS

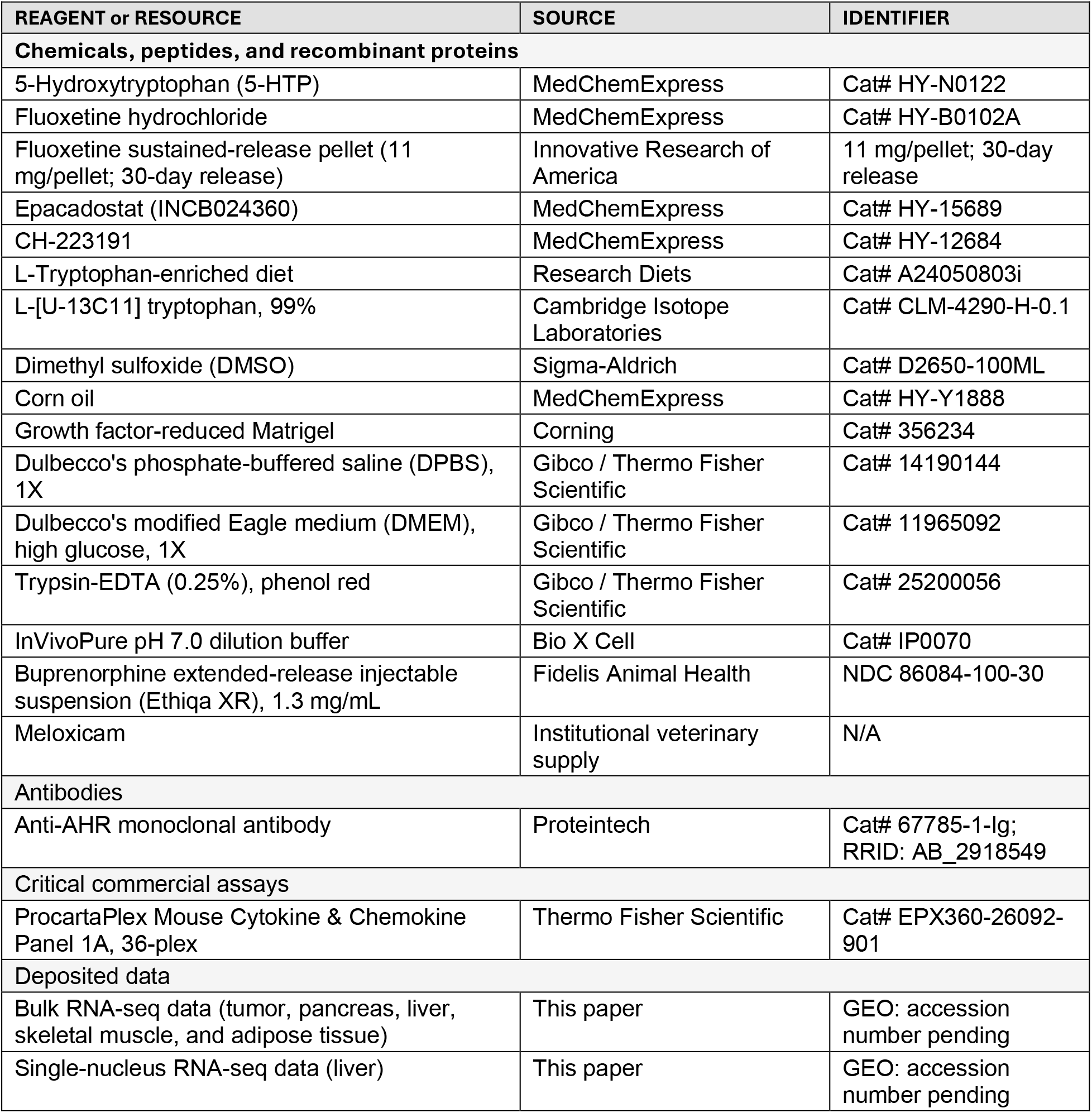
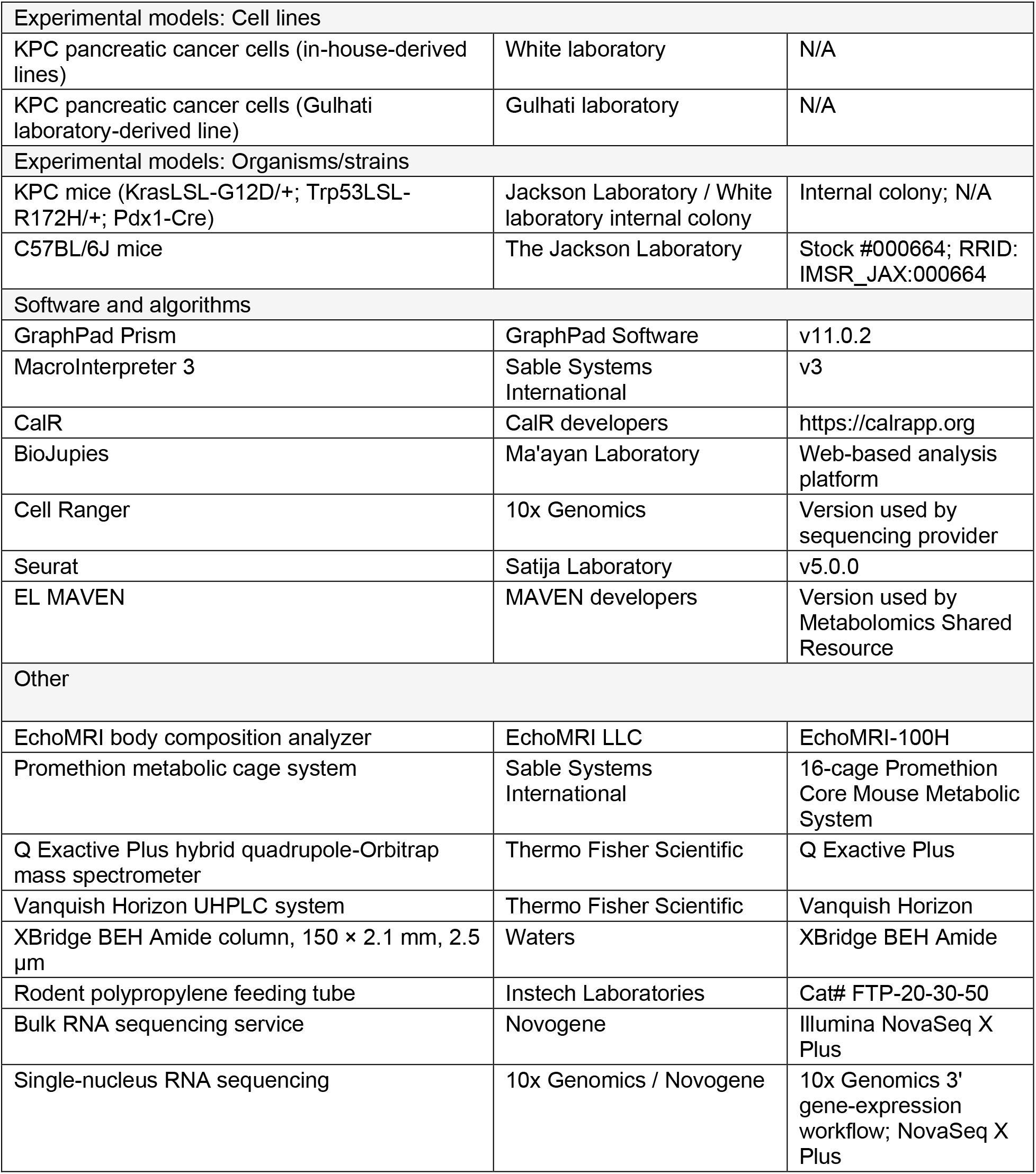

## RESOURCE AVAILABILITY

### Lead contact

Further information and requests for resources and reagents should be directed to and will be fulfilled by the lead contact, Eileen White.

### Materials availability

KPC pancreatic cancer cell lines generated or maintained in this study are available from the lead contact upon reasonable request and, where applicable, completion of an institutional material transfer agreement.

### Data and code availability

Bulk RNA-seq and single-nucleus RNA-seq data generated in this study will be deposited in the NCBI Gene Expression Omnibus (GEO). Accession numbers will be provided before final publication. This paper does not report original code. Any additional information required to reanalyze the data reported in this paper is available from the lead contact upon request.

### Mouse models

All animal studies were performed in accordance with protocols approved by the Rutgers University Institutional Animal Care and Use Committee (IACUC). Mice were maintained under specific pathogen-free conditions in a temperature- and humidity-controlled facility under a 12 h light/12 h dark cycle with ad libitum access to food and water unless otherwise indicated.

KPC mice carrying conditional KrasLSL-G12D/+ and Trp53LSL-R172H/+ alleles together with Pdx1-Cre were maintained as an internal colony. Both male and female mice were included across experiments. Littermate animals lacking Pdx1-Cre-mediated recombination served as non-tumor bearing controls and did not develop pancreatic tumors. For longitudinal studies, mice were monitored throughout tumor progression, with body weight and food intake measured at regular intervals and animals followed until prespecified humane endpoint criteria were reached. Cachexia status was defined relative to baseline body weight, as specified for individual experiments. Both male and female mice were included across experiments. KPC mice were approximately 4–5 months old at the cachectic stage. For orthotopic PDAC studies, 3-month-old C57BL/6J mice were obtained from The Jackson Laboratory and reached approximately 4 months of age by the cachectic stage.

C57BL/6J mice (The Jackson Laboratory, Stock #000664) were used as syngeneic recipients for orthotopic pancreatic tumor implantation experiments.

### KP and KL lung cancer models

KP (Kras^LSL-G12D/+;^ Trp53^fl/fl^) and KL (Kras^LSL-G12D/+^; Lkb1^fl/fl^) mice were generated and maintained in house. Both male and female mice were used. Lung tumors were initiated at 8–10 weeks of age by intranasal administration of adenovirus-Cre at 4 × 107 plaque-forming units (PFU) per mouse.

Following tumor initiation, mice were monitored longitudinally for disease progression and body weight loss and were analyzed at the cachectic stage, corresponding to approximately 15–20% loss of body weight. The KP model corresponds to the strain available from The Jackson Laboratory (RRID: IMSR_JAX:032435).

Individual experiments were conducted using male mice, female mice, or both sexes, depending on the experimental cohort. When both sexes were included within an experiment, data were pooled for analysis unless otherwise indicated. The study was not designed or powered to systematically assess sex-dependent differences.

### KPC derived pancreatic cancer cell lines

Four independent KPC tumor-derived pancreatic cancer cell lines were used to ensure reproducibility across biologically distinct tumor backgrounds. Two lines were established from female KPC tumors, including one line obtained from the Gulhati laboratory and one generated in house. Two additional independent lines were generated from male KPC tumors. For in house line generation, pancreatic tumors were harvested from tumor bearing mice, mechanically dissociated, and enzymatically digested to generate single cell suspensions. Cells were plated for primary outgrowth and serially passaged to enrich stable tumor cell populations and reduce stromal contamination. Cells were maintained in high glucose DMEM supplemented with 10% fetal bovine serum and 1% penicillin-streptomycin at 37^°C^ in a humidified incubator containing 5% CO_2_.

## METHOD DETAILS

### Orthotopic pancreatic tumor implantation

Orthotopic tumors were generated by surgical implantation of KPC pancreatic cancer cells into the pancreas of C57BL/6J recipient mice. Mice were anesthetized with inhaled isoflurane delivered through a precision vaporizer, and all procedures were performed under aseptic conditions. Perioperative analgesia consisted of meloxicam and extended-release buprenorphine (Ethiqa XR; 1.3 mg/mL; approximately 2.17 mg/kg) according to the approved Rutgers IACUC protocol. After shaving and preparation of the left abdominal surgical area, a small left-flank incision was made to expose the pancreas. A total of 2 × 10^5^ pancreatic cancer cells suspended in 10 μL growth factor-reduced Matrigel were injected into the pancreatic tail. The pancreas was gently returned to the abdominal cavity, and the incision was closed with absorbable sutures and wound clips. Mice were maintained on a warming pad throughout surgery and monitored continuously until full recovery from anesthesia. Animals were monitored postoperatively for 3 days according to IACUC guidelines and subsequently assessed daily for body weight, general health, and signs of distress. Wound clips were removed 14 days after surgery.

### Body composition measurements

Body composition was assessed longitudinally using an EchoMRI-100H body composition analyzer. Before each acquisition session, mice were weighed on a calibrated laboratory balance and instrument performance was verified using the manufacturer-recommended calibration standard. Fat mass, lean mass, free water, and total body water were measured in conscious, non-anesthetized mice. Animals were placed individually into restraint tubes and analyzed for approximately 2 min per mouse before being returned to their home cages. Measurements were collected at baseline and at designated time points during tumor progression.

### Ultrasound imaging

In vivo ultrasound imaging was performed using a Vevo 3100 high-frequency preclinical ultrasound system (FUJIFILM VisualSonics, Toronto, Canada). Mice were positioned on the imaging platform, and ultrasound gel was applied to the region of interest to ensure acoustic coupling. Images were acquired in B-mode using consistent acquisition settings across animals and experimental groups. Tumor dimensions were measured from ultrasound images and used to assess tumor growth.

### Metabolic cage analysis

Whole-body energy metabolism, feeding, and activity were assessed using a 16-cage Promethion Core Mouse Metabolic System (Sable Systems International). Mice were maintained on standard chow and singly housed for 48–72 h before the start of metabolic monitoring. During data collection, animals remained singly housed for 7 days under a 12 h light/12 h dark cycle at 21^°C^ and 55% relative humidity. The first 24 h of recording were excluded as an acclimation period. Oxygen consumption (VO_2_), carbon dioxide production (VCO_2_), food intake, energy expenditure, voluntary wheel running, and locomotor activity were recorded at 15 min intervals. Respiratory exchange ratio was calculated as VCO_2_/VO_2_. Horizontal and vertical activity were quantified using X-, Y-, and Z-axis infrared beam breaks. Stationary activity was defined as repeated interruption of a single beam, whereas ambulatory movement was defined by sequential interruption of two or more distinct beams. Raw Promethion files were processed using MacroInterpreter 3 and standardized outputs were subsequently analyzed using CalR (https://calrapp.org).

### Cage occupancy density maps and home-cage video recording

Raw metabolic-cage data were processed in R using the SableBase package (version 1.0.0) to read Sable Systems .exp files. X–Y position data were visualized using the R smoothScatter() function with nbin = 128, corresponding to a 128 × 128 grid. For group-level maps, X and Y position values were averaged across mice within each experimental group using rowMeans(), and the resulting group-level coordinates were used to generate the density maps. Videos S1 and S2 were recorded by camera during behavioral assessment.

### Cytokine profiling by multiplex Luminex assay

Cytokine profiling was performed using the ProcartaPlex Mouse Cytokine & Chemokine Panel 1A, 36-plex magnetic bead-based immunoassay. Following euthanasia, whole blood was collected by terminal cardiac puncture and allowed to clot at room temperature. Serum was isolated by centrifugation and stored at −80^°C^ until analysis. Samples were thawed on ice and centrifuged to remove residual particulate material. A total of 25 μL serum per sample was submitted to the Rutgers Cancer Institute of New Jersey Immune Monitoring and Flow Cytometry Shared Resource. Assay processing, acquisition, quality control, and analyte quantification were performed by the core facility according to manufacturer recommendations and standardized core procedures. Cytokine concentrations were determined by standard curve-based quantification and are reported as pg/mL.

### Bulk RNA sequencing and transcriptomic analysis

For transcriptomic analyses, tissues were collected from pre-cachectic and cachectic pancreatic ductal adenocarcinoma (PDAC) mice, defined as animals exhibiting 5%–15% and >15% body weight loss from baseline, respectively. Tissues were rapidly dissected, flash-frozen in liquid nitrogen, and stored at −80^°C^ until processing. RNA extraction, sample quality control, mRNA library preparation, sequencing, and primary sequencing quality control were performed by Novogene. Total RNA was extracted from frozen tumor, pancreas, liver, skeletal muscle, and adipose tissues, and non-stranded mRNA libraries were generated following poly(A) enrichment. Libraries were sequenced on an Illumina NovaSeq X Plus platform using paired-end 150-bp reads with a target output of approximately 6 Gb raw sequencing data per sample. Adapter-containing and low-quality reads were removed during primary quality-control processing. Processed gene-expression datasets were analyzed using BioJupies for differential gene-expression and pathway-enrichment analyses. Differentially expressed genes were used for downstream pathway-level interpretation and visualization, and functional enrichment outputs generated by Novogene and BioJupies were used to identify biological processes and signaling pathways altered during PDAC cachexia progression.

For visualization of selected transcriptional programs, normalized expression values were imported into GraphPad Prism v11.0.2. Individual gene expression values were z-score normalized across all samples within each tissue, and heatmaps were generated from the resulting standardized expression matrices.

### Single-nucleus RNA sequencing and analysis

Single-nucleus RNA sequencing was performed on frozen liver samples from nine mice, comprising three controls, three GEMM KPC mice, and three orthotopic PDAC-bearing mice. Nuclei isolation and library preparation were performed by Novogene using their standard single-nucleus workflow. Nuclei were released by tissue lysis and mechanical dissociation, followed by washing, filtration, counting, and quality assessment before library preparation. Libraries were generated using the 10x Genomics Chromium Single Cell 3′ gene-expression workflow, with a target recovery of approximately 7,000 nuclei per sample, and sequenced on an Illumina NovaSeq X Plus platform using paired-end 150-bp reads at a target depth of approximately 50,000 read pairs per nucleus.

Raw reads were processed using Cell Ranger v7.0.0 (10x Genomics) with the mm10-2020-A mouse reference transcriptome for read alignment, cell-barcode and unique molecular identifier processing, and generation of gene-by-nucleus count matrices. Intronic reads were included in the count matrix. Count matrices were imported into Seurat v5.0.0.^93^ Nuclei were retained if they contained more than 250 detected features, more than 500 total RNA counts, and less than 10% mitochondrial transcripts. A total of 98,255 nuclei were retained across the nine samples. Each dataset was normalized using NormalizeData, and highly variable features were identified using FindVariableFeatures with the variance-stabilizing transformation method and 5,000 features. All nine datasets were integrated using canonical correlation analysis. SelectIntegrationFeatures was performed using 5,000 features, followed by FindIntegrationAnchors and IntegrateData. ScaleData, RunPCA, and RunUMAP were run on both integrated and unintegrated data using 30 principal components. Louvain clustering was performed using 30 principal components for FindNeighbors and a resolution of 0.05 for FindClusters. Marker genes for each cluster were identified using FindAllMarkers with default parameters. Cluster identities were assigned using a published mouse liver reference atlas^94^ by determining the reference cell type in which each cluster’s top five marker genes showed the highest expression. Overall, ten liver cell types were identified among eleven clusters.

### Drug administration by oral gavage and intraperitoneal injection

Experimental compounds were administered by oral gavage or intraperitoneal (i.p.) injection according to the individual study design. Oral gavage was performed using polypropylene rodent feeding tubes (Instech Laboratories, Cat# FTP-20-30-50). Mice were gently restrained, and dosing volumes were adjusted according to compound concentration, experimental dose, and body weight. For i.p. administration, compounds were prepared in the indicated vehicle and administered at 100–200 μL per mouse. Treatment-specific formulations and schedules are described below.

### Fluoxetine pellet administration

Sustained-release fluoxetine pellets (Innovative Research of America; 11 mg/pellet, 30-day release) were implanted subcutaneously under isoflurane anesthesia, corresponding to an estimated release rate of approximately 0.37 mg/day (approximately 15 mg/kg/day). A small incision was made in the dorsal interscapular region and a subcutaneous pocket was created for pellet placement. Meloxicam was administered for perioperative analgesia, and incisions were closed with surgical staples. Procedures were performed on a heating pad to maintain body temperature. Animals were continuously monitored following implantation and returned to their home cages after full recovery from anesthesia.

### IDO1 inhibition

Tumor bearing mice were treated with the selective IDO1 inhibitor Epacadostat (INCB024360; MedChemExpress, Cat# HY-15689). Treatment was initiated 4 days after orthotopic tumor implantation. Epacadostat was first dissolved in DMSO and then diluted in corn oil to a final vehicle composition of 10% DMSO and 90% corn oil. Mice received Epacadostat at 100 mg/kg by oral gavage once daily until the experimental endpoint. Vehicle-treated tumor bearing controls received an equivalent volume of 10% DMSO/90% corn oil on the same schedule.

### Fluoxetine treatment

Tumor bearing mice were treated with fluoxetine hydrochloride (MedChemExpress, Cat# HY-B0102A; CAS 56296-78-7) beginning 4 days after orthotopic tumor implantation. Fluoxetine was formulated at 3.75 mg/mL in 50% (v/v) PEG300 and 50% (v/v) sterile 0.9% saline and administered by intraperitoneal injection at 15 mg/kg once daily until the study endpoint. Control mice received an equivalent volume of vehicle.

### 5-HTP treatment

Tumor bearing mice were treated with 5-hydroxytryptophan (5-HTP; MedChemExpress, Cat# HY-N0122) by intraperitoneal injection at 100 mg/kg once daily until the study endpoint. 5-HTP was formulated in a vehicle consisting of 10% DMSO and 90% 20% SBE-β-CD in saline (v/v). Control mice received an equivalent volume of the corresponding vehicle.

### AhR inhibition

To evaluate the contribution of aryl hydrocarbon receptor (AhR) signaling during pancreatic cancer cachexia, tumor bearing mice were treated with the AhR antagonist CH-223191 (MedChemExpress, Cat# HY-12684). Treatment was initiated 4–5 days after orthotopic tumor implantation. CH-223191 was formulated in 10% DMSO and 90% corn oil and administered by oral gavage at 10 mg/kg/day until study endpoint. Control mice received vehicle alone on the same schedule.

### Combined fluoxetine and IDO1 inhibition

For combined pathway targeting, tumor bearing mice received fluoxetine and epacadostat beginning 5 days after orthotopic tumor implantation. Fluoxetine was delivered using a sustained-release subcutaneous pellet (11 mg/pellet, 30-day release), corresponding to an estimated dose of approximately 15 mg/kg/day for a 25 g mouse. Epacadostat was administered by oral gavage at 100 mg/kg/day in 10% DMSO/90% corn oil. Combination treatment was maintained until the experimental endpoint. Control mice received placebo pellets together with daily vehicle gavage.

### High-tryptophan diet

For dietary tryptophan supplementation experiments, orthotopic PDAC-bearing mice were randomized to control or 10× tryptophan-enriched diets beginning 5 days after tumor implantation. Body composition was assessed by EchoMRI during disease progression. The dietary intervention was maintained until the experimental endpoint, with analyses performed on day 12 and at the cachectic endpoint. The custom tryptophan-enriched diet was obtained from Research Diets (Cat# A24050803i).

### Tumor collection and tissue processing

At experimental endpoint, mice were euthanized by CO_2_ inhalation followed by cervical dislocation. Blood was collected by cardiac puncture and serum isolated by centrifugation. Tumors, liver, spleen, skeletal muscle, and adipose tissues were rapidly harvested and weighed. Tumor samples designated for metabolic analysis were mechanically compressed using a pre-chilled tissue clamp, flash-frozen in liquid nitrogen, and transferred to −80°C for long-term storage. Tissues designated for metabolomic or transcriptomic analyses were maintained at −80°C until processing.

### Metabolomics analysis by LC-MS

Tissues were rapidly excised, freeze-clamped in liquid nitrogen using a pre-chilled Wollenberger clamp, and stored at −80^°C^ until processing. Frozen tissue (10–30 mg) was transferred to pre-labeled tubes containing grinding beads and mechanically pulverized under cryogenic conditions on dry ice. Polar metabolites were extracted using methanol:acetonitrile:water (40:40:20, v/v/v) containing 0.5% formic acid, followed by neutralization with ammonium bicarbonate. After sequential extraction and centrifugation, clarified supernatants were collected, supplemented with internal standard, and stored at −80^°C^ until analysis.

LC-MS analysis was performed by the Rutgers Cancer Institute Metabolomics Shared Resource using a Q Exactive Plus hybrid quadrupole-Orbitrap mass spectrometer coupled to a Vanquish Horizon UHPLC system (Thermo Fisher Scientific). Metabolites were separated on an XBridge BEH Amide column (150 × 2.1 mm, 2.5 μm; Waters) under hydrophilic interaction liquid chromatography conditions using ammonium hydroxide- and ammonium acetate-containing mobile phases. Data were acquired in negative ionization mode over m/z 72–1000 at a resolution of 70,000 at m/z 200. Metabolites were identified and quantified using El-MAVEN v0.12.1-beta with a mass-accuracy threshold of 5 ppm. Metabolite ion counts were expressed as fold change relative to the control mean and log10-transformed for statistical analysis. Undetected metabolites were treated as missing; therefore, n varied by metabolite. Heatmaps included samples with complete metabolite profiles, whereas bar plots included all samples in which the corresponding metabolite was detected.

### Histology and immunohistochemistry

Pancreatic tumors from IDO1 inhibitor-treated and vehicle-treated mice were collected at humane endpoint, fixed overnight in 10% neutral-buffered formalin, transferred to 70% ethanol, and paraffin embedded. Sections were deparaffinized and rehydrated. Hematoxylin and eosin staining was performed for histopathologic assessment. For immunohistochemistry, antigen retrieval was performed in 10 mM sodium citrate buffer (pH 6.0), and sections were incubated with anti-AHR monoclonal antibody (Proteintech, Cat# 67785-1-Ig; RRID: AB_2918549) at 1:400 dilution. The specificity of the AhR antibody was supported by prior validation in published studies and by the expected nuclear/cytoplasmic staining pattern observed in tumor tissue. Images were acquired using a Nikon Eclipse 80i microscope. Three animals per experimental group were included in the histologic analyses. Nuclear AhR positivity was quantified using QuPath (v0.5; Bankhead et al., 2017). Stain vectors were estimated from representative images, and nuclei were detected using the Positive cell detection command. Nuclei with a mean DAB optical density above 0.6 were classified as AhR-positive. Identical parameters were applied to all images, and detections were visually verified to exclude erythrocytes and necrotic regions. Results are expressed as the percentage of AhR-positive nuclei.

### Forced swim test

Immobility behavior was assessed using the forced swim test. Mice were individually placed in transparent cylindrical containers approximately 20–25 cm high and 10–15 cm in diameter, filled with water maintained at 23–25^°C^ to a depth that prevented tail support while allowing unrestricted swimming. Each animal underwent a 6 min session. The first 1 min was treated as an acclimation period and excluded from analysis; behavior was quantified during the final 5 min. Scoring was performed blinded to experimental group. Immobility was defined as the absence of active escape directed movements, with only movements required to maintain flotation and keep the head above water. Mobility time (total test time minus immobility time) and latency to the first immobility episode were quantified. After testing, mice were removed from the water, gently dried, allowed to recover under warming conditions, and returned to their home cages.

### In vivo stable isotope tracing

Mice bearing chronic jugular vein catheters received a continuous infusion of L-U-¹³C₁₁-tryptophan (99%; Cambridge Isotope Laboratories, Cat# CLM-4290-H-0.1), prepared in sterile 0.9% saline at 5 mg/mL (approximately 23 mM). The tracer was infused at 0.1 μL/g body weight/min, corresponding to approximately 2.3 nmol/g/min, for 2.5 h. At the end of the infusion, mice were euthanized and serum, pancreas/tumor, and liver were rapidly collected. Tissues were immediately freeze-clamped using pre-chilled Wollenberger clamps, snap frozen in liquid nitrogen, and stored at −80°C until metabolite extraction. Labeling of tryptophan and downstream kynurenine, serotonin, and indole pathway metabolites was quantified by LC-MS. Isotopologue distributions were corrected for natural isotope abundance and normalized to serum tryptophan enrichment before downstream analysis.

### Adenylate energy charge calculation

Adenylate energy charge was calculated for soleus muscle, liver, and spleen from LC-MS measurements of ATP, ADP, and AMP using the formula: (ATP + 0.5 × ADP)/(ATP + ADP + AMP). Energy charge values were calculated independently for each sample and used for statistical comparisons across experimental groups.

## QUANTIFICATION AND STATISTICAL ANALYSIS

Statistical analyses were performed using GraphPad Prism v11.0.2. Data are presented as mean ± SEM unless otherwise indicated. Comparisons between two groups were performed using unpaired two-tailed Student’s t tests. Comparisons involving multiple groups were performed using one-way or two-way ANOVA, as appropriate. Longitudinal and metabolic cage data were analyzed by two-way repeated-measures ANOVA or, when values were missing, a mixed-effects model with Geisser–Greenhouse correction; for metabolic cage data, the group effect is reported. Multiple comparisons were performed using Tukey’s, Dunnett’s, or Šídák’s tests or uncorrected Fisher’s LSD test, as indicated in the figure legends. Metabolite fold changes were log10-transformed before analysis. Survival was analyzed by the log-rank test. No statistical methods were used to predetermine sample sizes. For in vivo experiments, n denotes individual animals unless otherwise stated. Exact sample sizes, statistical tests, and p values are provided in the corresponding figure legends. Differences were considered statistically significant at p < 0.05.

---

**Figure S1.**
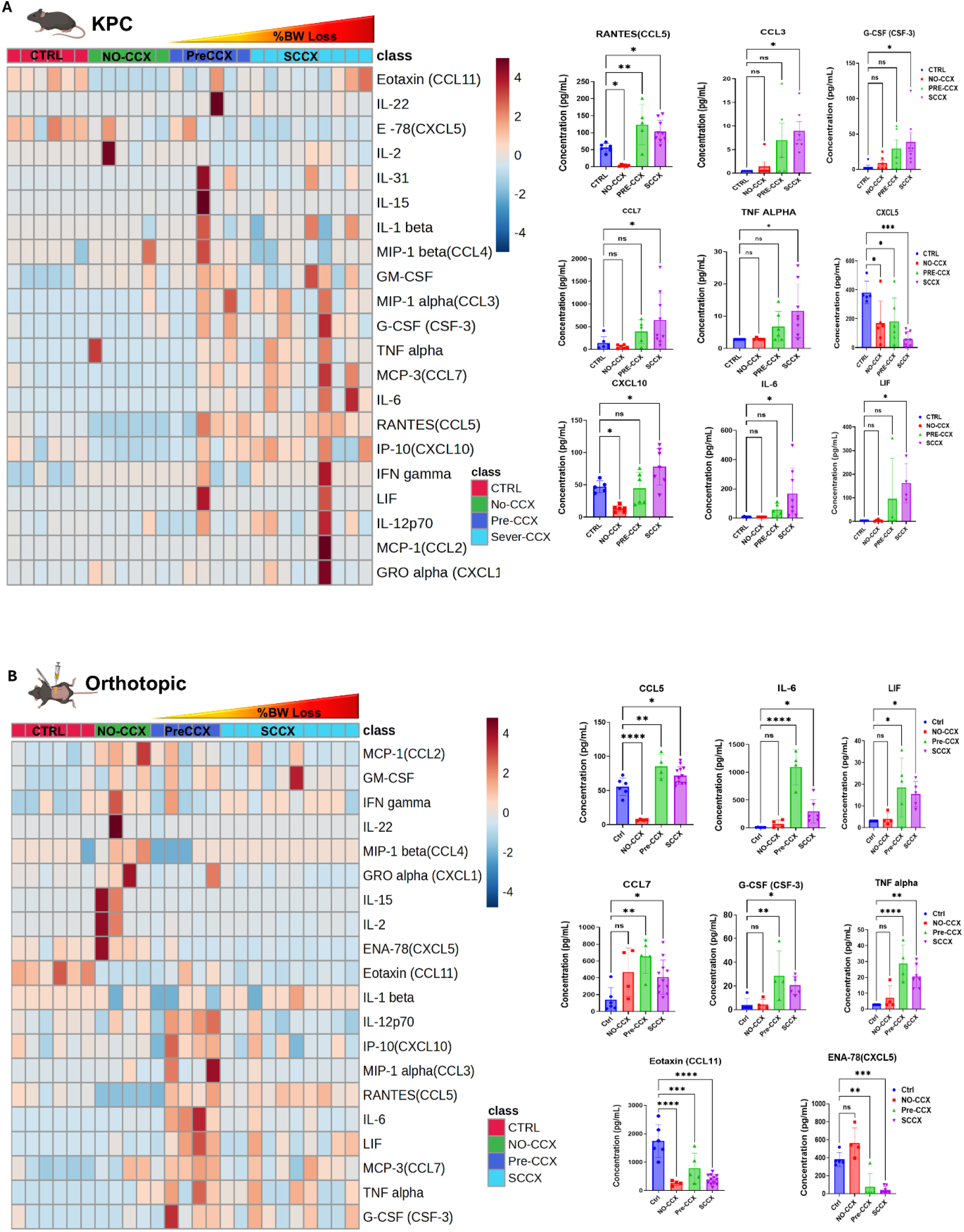
Circulating inflammatory profiles across stages of PDAC cachexia, related to Figure 1. (A) Circulating cytokine and chemokine profiles in control (CTRL; n = 6), non-cachectic (NO-CCX; n = 6), pre-cachectic (PreCCX; n = 6), and severe cachexia (SCCX; n = 9) GEMM KPC mice, shown as a heatmap with selected analytes. (B) Corresponding profiles in orthotopic PDAC mice: CTRL (n = 6), NO-CCX (n = 4), PreCCX (n = 5), and SCCX (n = 13). NO-CCX, non-cachectic; PreCCX, pre-cachectic; SCCX, severe cachexia; CCX, cachexia. Data are mean ± SEM. Individual analytes were analyzed by ordinary one-way ANOVA with Dunnett’s multiple-comparisons test versus CTRL. Undetected analytes were excluded; therefore, n varies by analyte. *p < 0.05, **p < 0.01, ***p < 0.001, ****p < 0.0001; ns, p > 0.05.

**Figure S2.**
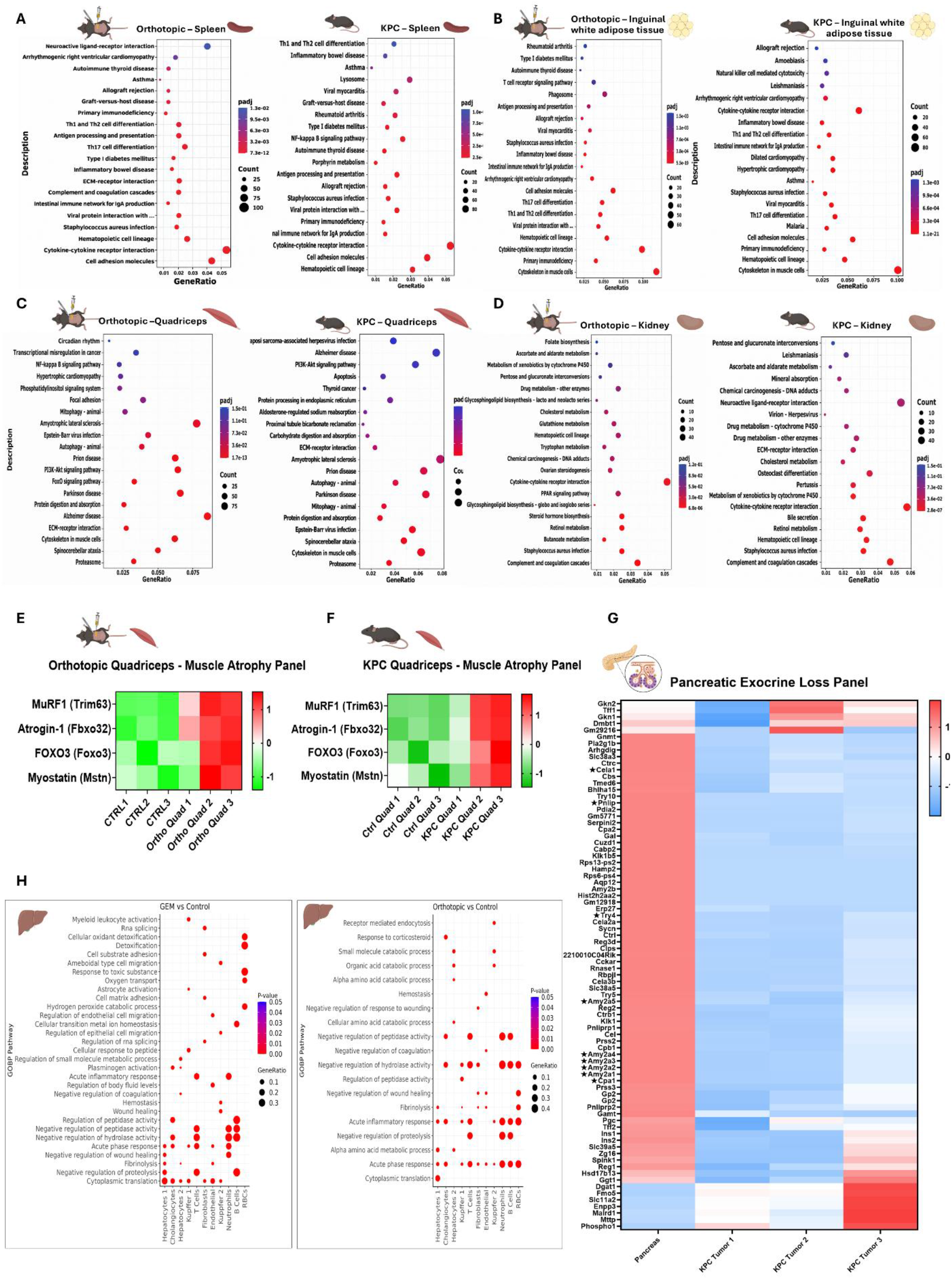
Tissue transcriptional changes across PDAC cachexia models, related to Figure 2. (A–D) Pathway enrichment of bulk RNA-seq profiles from orthotopic PDAC and GEMM KPC mice versus corresponding controls in spleen (A), inguinal white adipose tissue (B), quadriceps muscle (C), and kidney (D) (n = 3 mice per group). Dot size indicates gene count; color indicates adjusted p value. (E and F) Expression of selected muscle atrophy-associated genes in quadriceps from orthotopic PDAC-bearing (E) and GEMM KPC (F) mice versus controls (n = 3 mice per group). (G) Heatmap of pancreatic exocrine-associated gene expression across control and PDAC samples. (H) Cell type-resolved Gene Ontology enrichment from liver single-nucleus RNA-seq in GEMM KPC and orthotopic PDAC-bearing mice versus corresponding controls (n = 3 mice per group).

**Figure S3.**
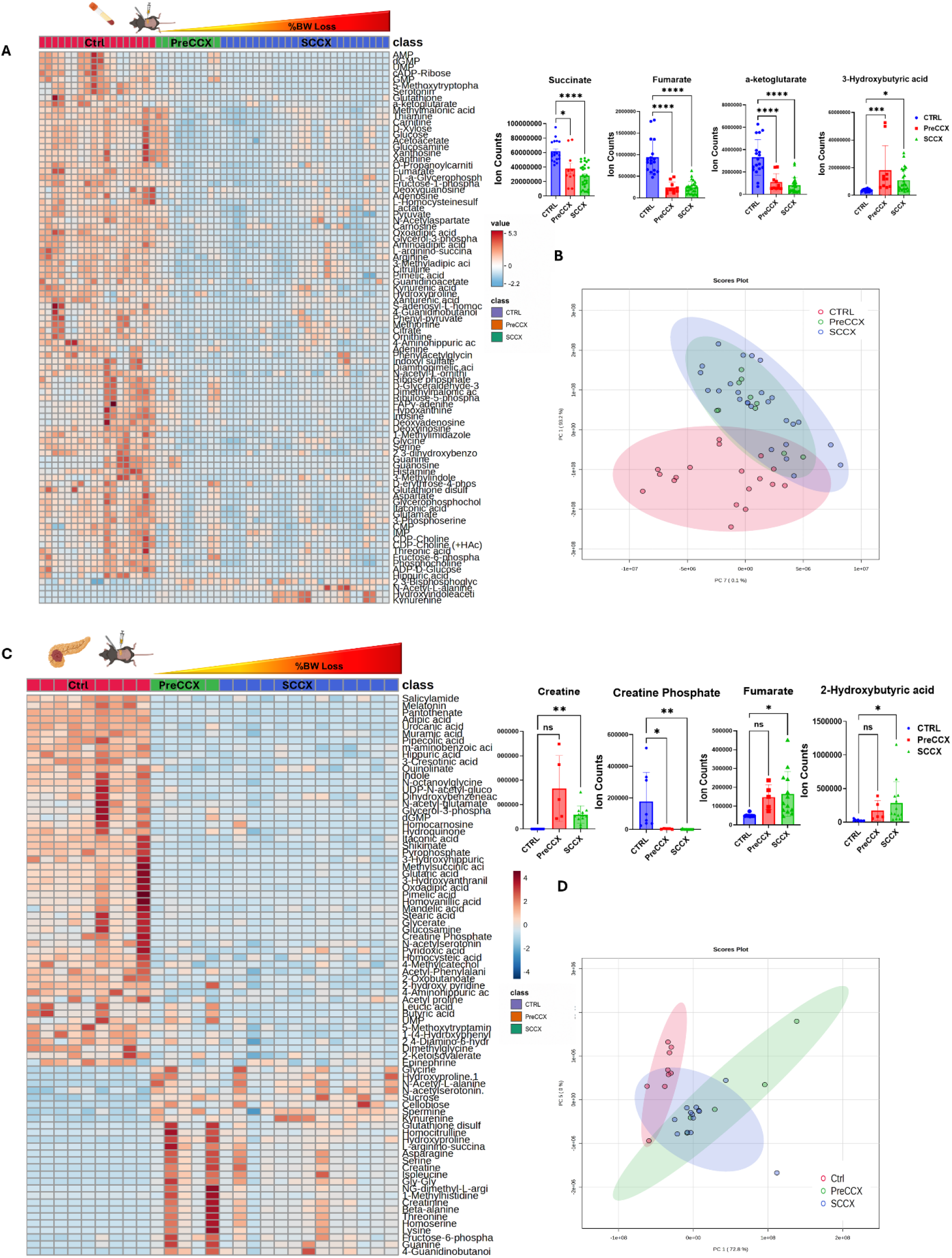
Metabolomic profiles during orthotopic PDAC cachexia progression, related to Figure 3. (A) Serum metabolite heatmap from control (CTRL; n = 18), pre-cachectic (PreCCX; n = 10), and severe cachexia (SCCX; n = 26) mice, with selected metabolites shown individually. (B) Principal-component analysis (PCA) of serum metabolomic profiles. (C) Pancreas/tumor metabolite heatmap from CTRL (n = 9), PreCCX (n = 5), and SCCX (n = 13) mice, with selected metabolites shown individually. (D) PCA of pancreas/tumor metabolomic profiles. PreCCX, pre-cachectic; SCCX, severe cachexia; CCX, cachexia. Data are mean ± SEM where applicable. Selected metabolites were analyzed by ordinary one-way ANOVA with Dunnett’s multiple-comparisons test versus CTRL. Undetected metabolites were excluded; therefore, n varies by metabolite. Heatmaps include samples with complete metabolomic profiles; Metabolites not detected in a given sample were excluded from analysis; therefore, n varies by metabolite bar plots include all samples in which the indicated metabolite was measured. *p < 0.05, **p < 0.01, ***p < 0.001, ****p < 0.0001; ns, p > 0.05.

**Figure S4.**
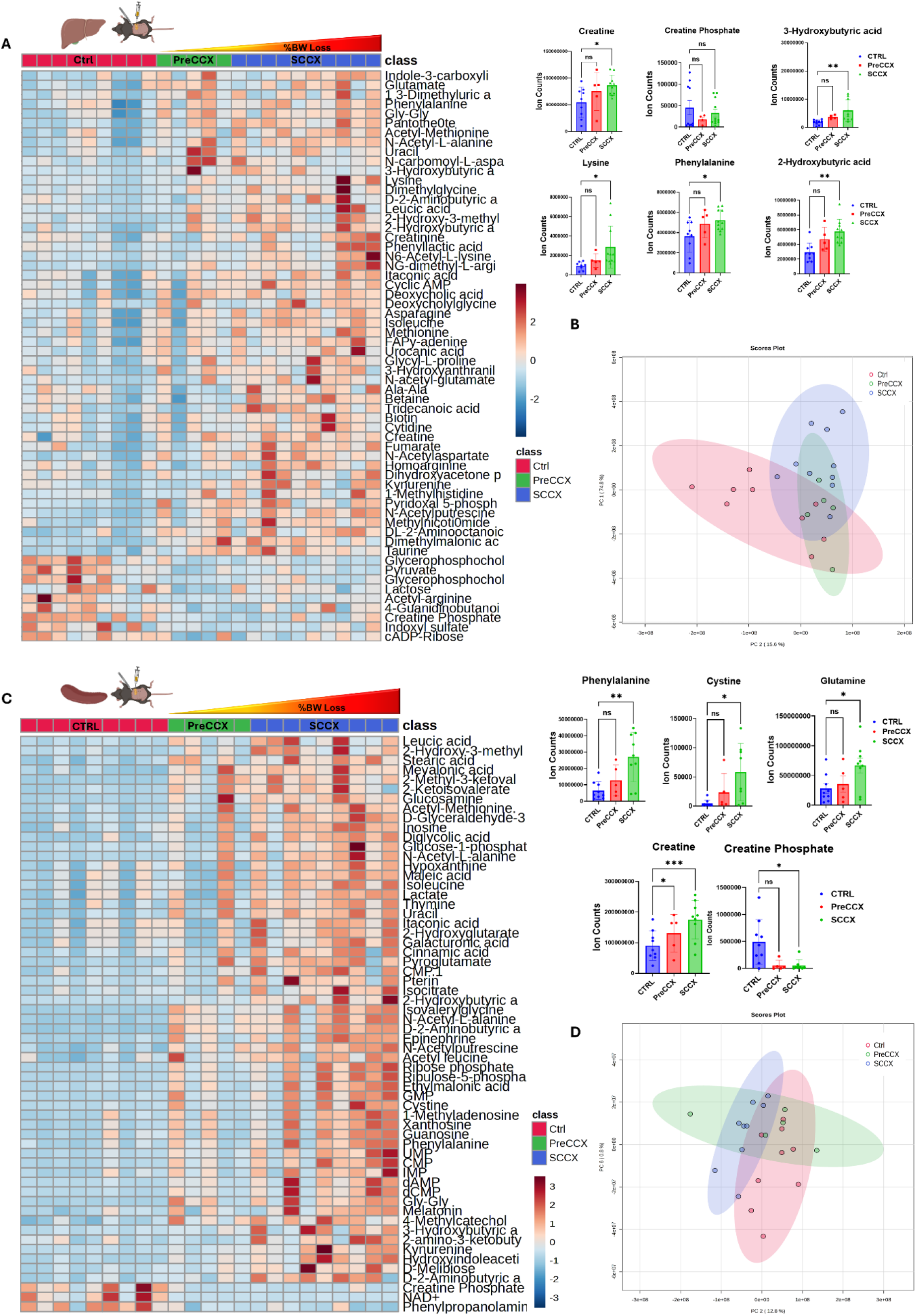
Liver and spleen metabolomic profiles during orthotopic PDAC cachexia progression, related to Figure 3. (A) Liver metabolite heatmap from control (CTRL; n = 9), pre-cachectic (PreCCX; n = 5), and severe cachexia (SCCX; n = 9) mice, with selected metabolites shown individually. (B) Principal-component analysis (PCA) of liver metabolomic profiles. (C) Spleen metabolite heatmap from CTRL (n = 9), PreCCX (n = 5), and SCCX (n = 10) mice, with selected metabolites shown individually. (D) PCA of spleen metabolomic profiles. PreCCX, pre-cachectic; SCCX, severe cachexia; CCX, cachexia. Data are mean ± SEM where applicable. Selected metabolites were analyzed by ordinary one-way ANOVA with Dunnett’s multiple-comparisons test versus CTRL. Undetected metabolites were excluded; therefore, n varies by metabolite. Heatmaps include samples with complete metabolomic profiles; Metabolites not detected in a given sample were excluded from analysis; therefore, n varies by metabolite bar plots include all samples in which the indicated metabolite was measured. *p < 0.05, **p < 0.01, ***p < 0.001, ****p < 0.0001; ns, p > 0.05.

**Figure S5.**
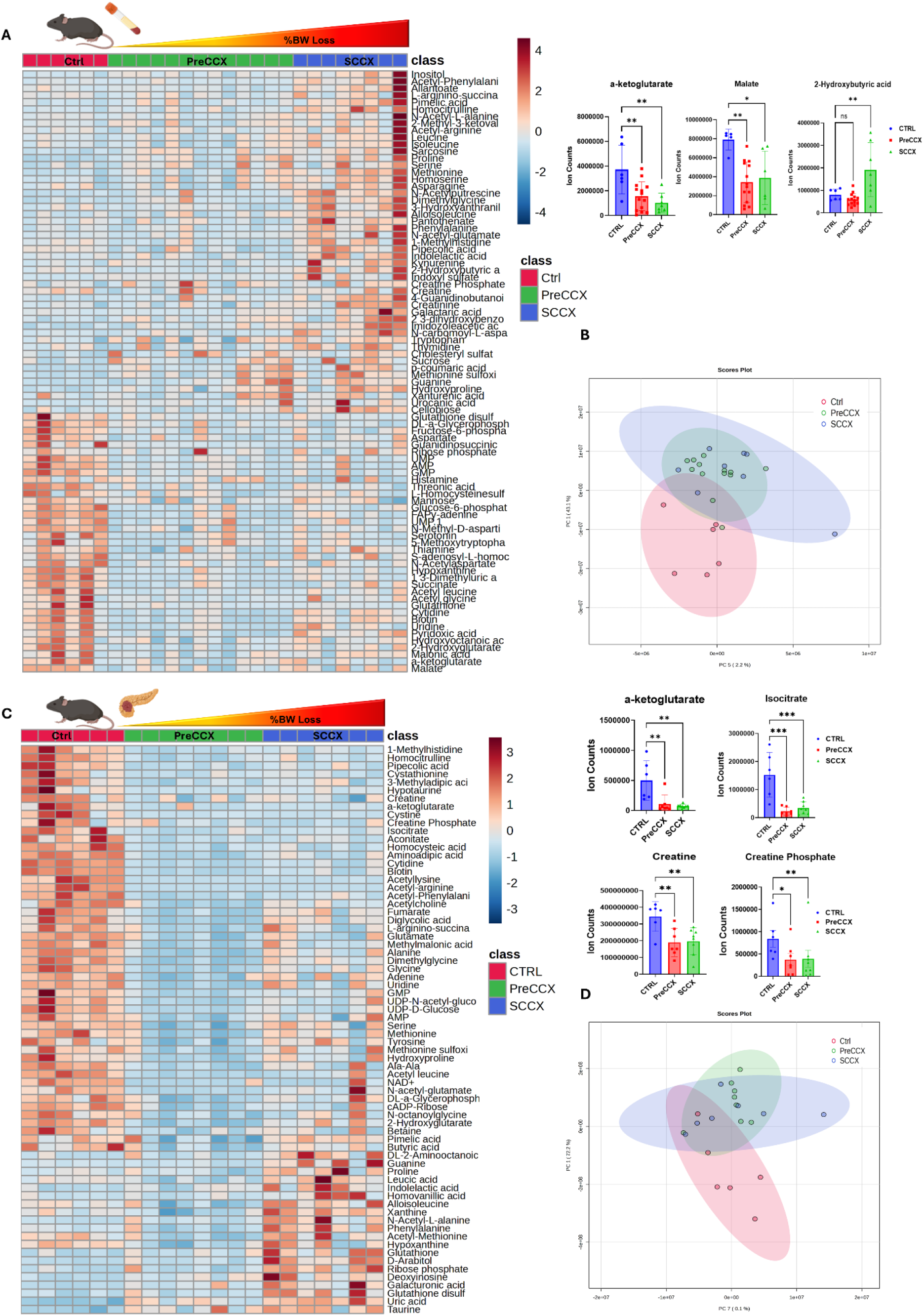
Circulating and pancreatic/tumor metabolic remodeling during GEMM KPC cachexia progression, related to Figure 3. (A) Serum metabolite heatmap from control (CTRL; n = 6), pre-cachectic (PreCCX; n = 13), and severe cachexia (SCCX; n = 8) GEMM KPC mice, with selected metabolites shown individually. (B) Principal-component analysis (PCA) of serum metabolomic profiles. (C) Pancreas/tumor metabolite heatmap from CTRL (n = 6), PreCCX (n = 8), and SCCX (n = 7) mice, with selected metabolites shown individually. (D) PCA of pancreas/tumor metabolomic profiles. PreCCX, pre-cachectic; SCCX, severe cachexia; CCX, cachexia. Data are mean ± SEM where applicable. Selected metabolites were analyzed by ordinary one-way ANOVA with Dunnett’s multiple-comparisons test versus CTRL. Undetected metabolites were excluded; therefore, n varies by metabolite. Heatmaps include samples with complete metabolomic profiles; Metabolites not detected in a given sample were excluded from analysis; therefore, n varies by metabolite bar plots include all samples in which the indicated metabolite was measured. *p < 0.05, **p < 0.01, ***p < 0.001, ****p < 0.0001; ns, p > 0.05.

**Figure S6.**
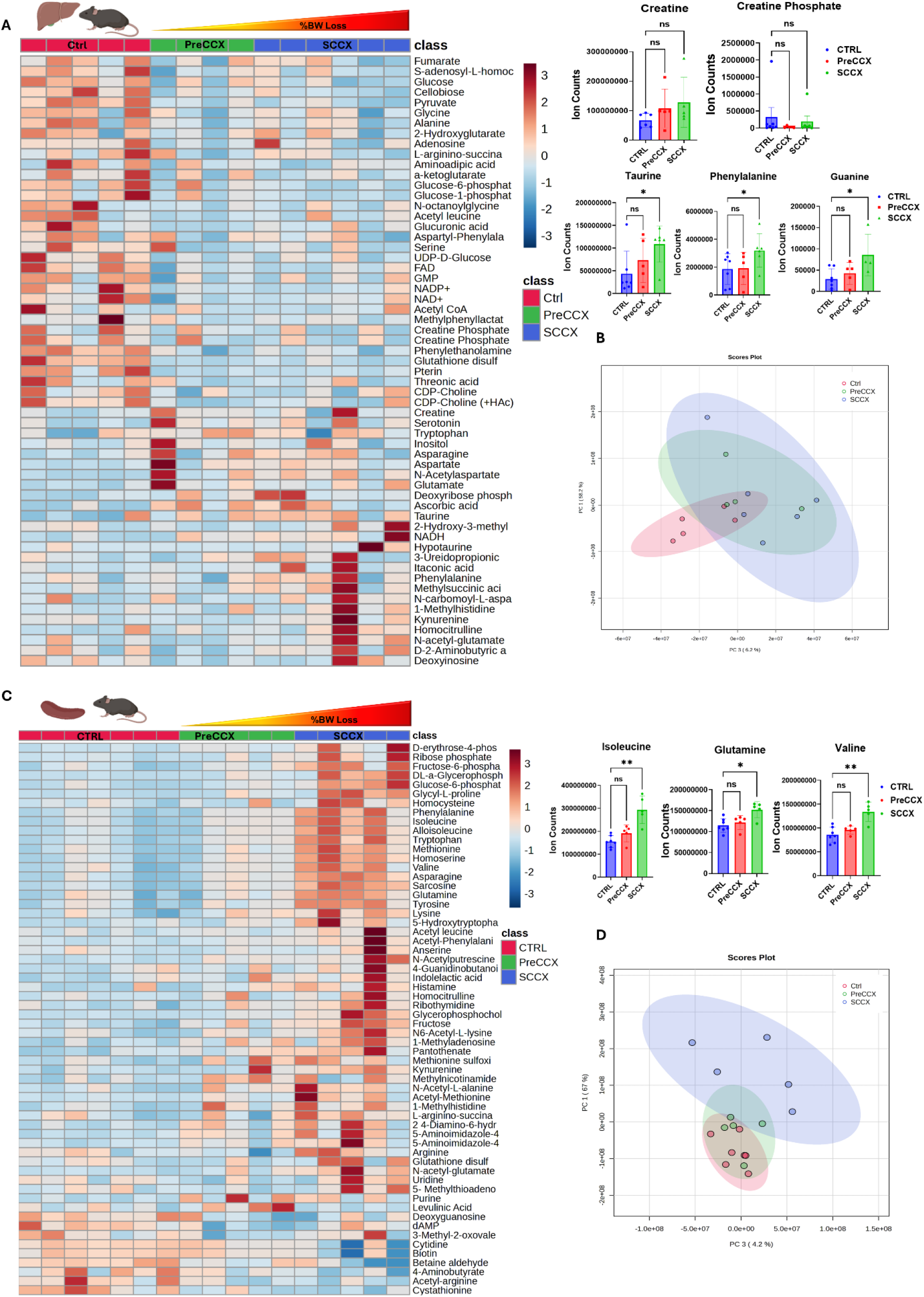
Hepatic and splenic metabolic remodeling during GEMM KPC cachexia progression, related to Figure 3. (A) Liver metabolite heatmap from control (CTRL; n = 5), pre-cachectic (PreCCX; n = 4), and severe cachexia (SCCX; n = 6) GEMM KPC mice, with selected metabolites shown individually. (B) Principal-component analysis (PCA) of liver metabolomic profiles. (C) Spleen metabolite heatmap from CTRL (n = 7), PreCCX (n = 5), and SCCX (n = 5) mice, with selected metabolites shown individually. (D) PCA of spleen metabolomic profiles. PreCCX, pre-cachectic; SCCX, severe cachexia; CCX, cachexia. Data are mean ± SEM where applicable. Selected metabolites were analyzed by ordinary one-way ANOVA with Dunnett’s multiple-comparisons test versus CTRL. Undetected metabolites were excluded; therefore, n varies by metabolite. Heatmaps include samples with complete metabolomic profiles; Metabolites not detected in a given sample were excluded from analysis; therefore, n varies by metabolite bar plots include all samples in which the indicated metabolite was measured. *p < 0.05, **p < 0.01, ***p < 0.001, ****p < 0.0001; ns, p > 0.05.

**Figure S7.**
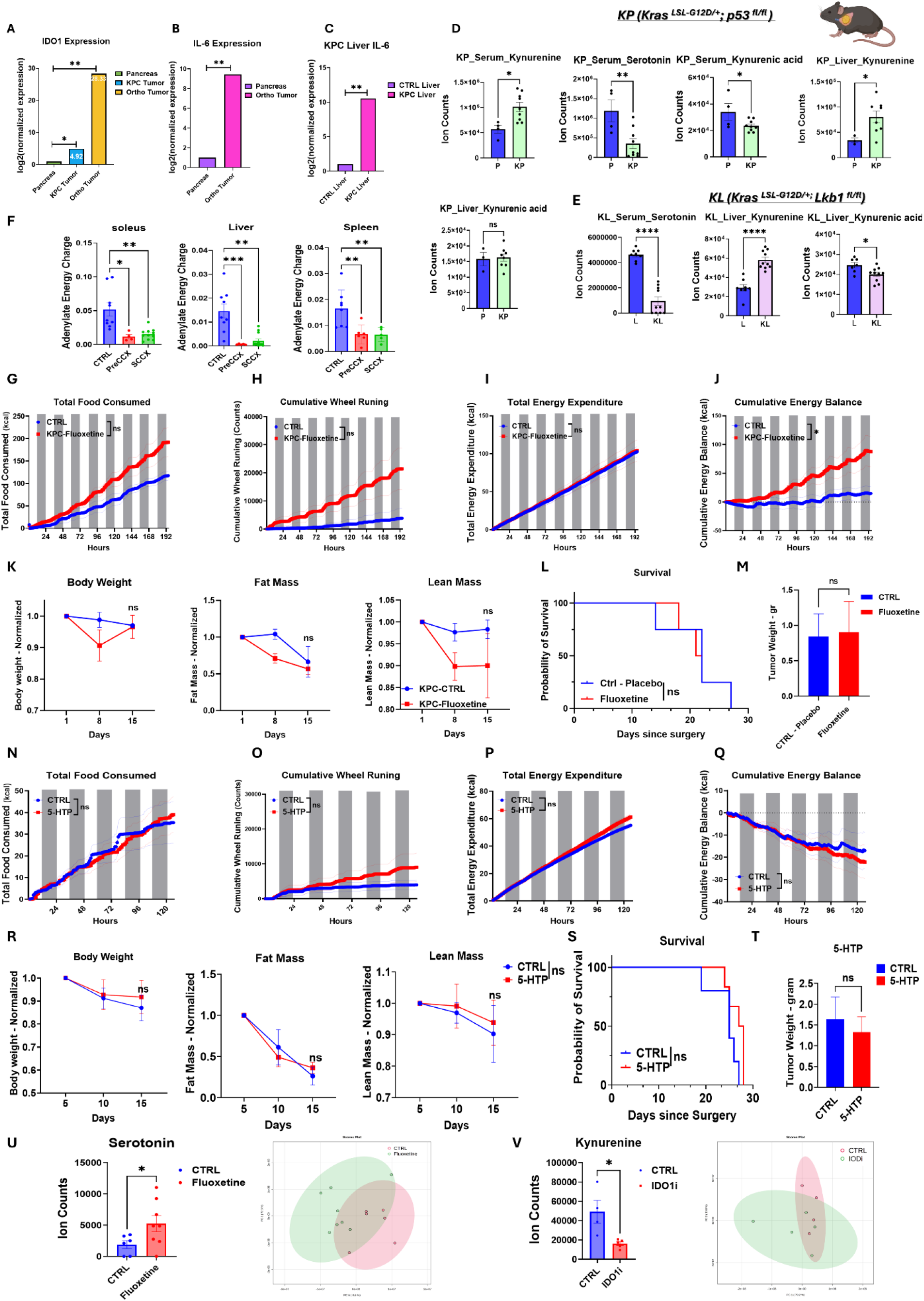
Tryptophan pathway and metabolic responses across cachexia models, related to Figures 4–6. (A–C) IDO1 and IL-6 expression in pancreas/tumor and liver as indicated. (D and E) Serum and liver tryptophan-pathway metabolites in KP (D) and KL (E) lung cancer models relative to controls. (F) Adenylate energy charge in soleus, liver, and spleen across CTRL, PreCCX, and SCCX stages. (G–J) Metabolic-cage analysis of mixed-sex GEMM KPC mice treated with placebo (n = 3) or sustained-release fluoxetine (n = 3): cumulative food intake (G), wheel running (H), energy expenditure (I), and energy balance (J). (K–M) Longitudinal body weight, fat mass, and lean mass (K), survival (L), and endpoint tumor weight (M) after fluoxetine treatment. (N–Q) Metabolic-cage analysis of mixed-sex orthotopic PDAC-bearing mice treated with vehicle (n = 5) or 5-HTP (n = 5): cumulative food intake (N), wheel running (O), energy expenditure (P), and energy balance (Q). (R–T) Longitudinal body weight, fat mass, and lean mass (R), survival (S), and endpoint tumor weight (T) after 5-HTP treatment. (U) Circulating serotonin and serum metabolomic PCA in control (n = 6) and fluoxetine-treated (n = 8) orthotopic PDAC-bearing mice. (V) Circulating kynurenine and serum metabolomic PCA in control (n = 4) and IDO1 inhibitor-treated (n = 5) orthotopic PDAC-bearing mice. PreCCX, pre-cachectic; SCCX, severe cachexia. Gray bars indicate dark phases; shading denotes SEM. Data are mean ± SEM. Metabolic-cage data were analyzed by two-way repeated-measures ANOVA or mixed-effects models with Geisser–Greenhouse correction followed by Šídák’s multiple-comparisons test. Statistics for other panels are indicated in the panels and STAR Methods. *p < 0.05, **p < 0.01, ***p < 0.001, ****p < 0.0001; ns, p > 0.05.

