## Supplemental Videos for "Behavioral dysfunction but not wasting in cancer cachexia is driven by altered tryptophan metabolism": videos.pptx

### Slide 1
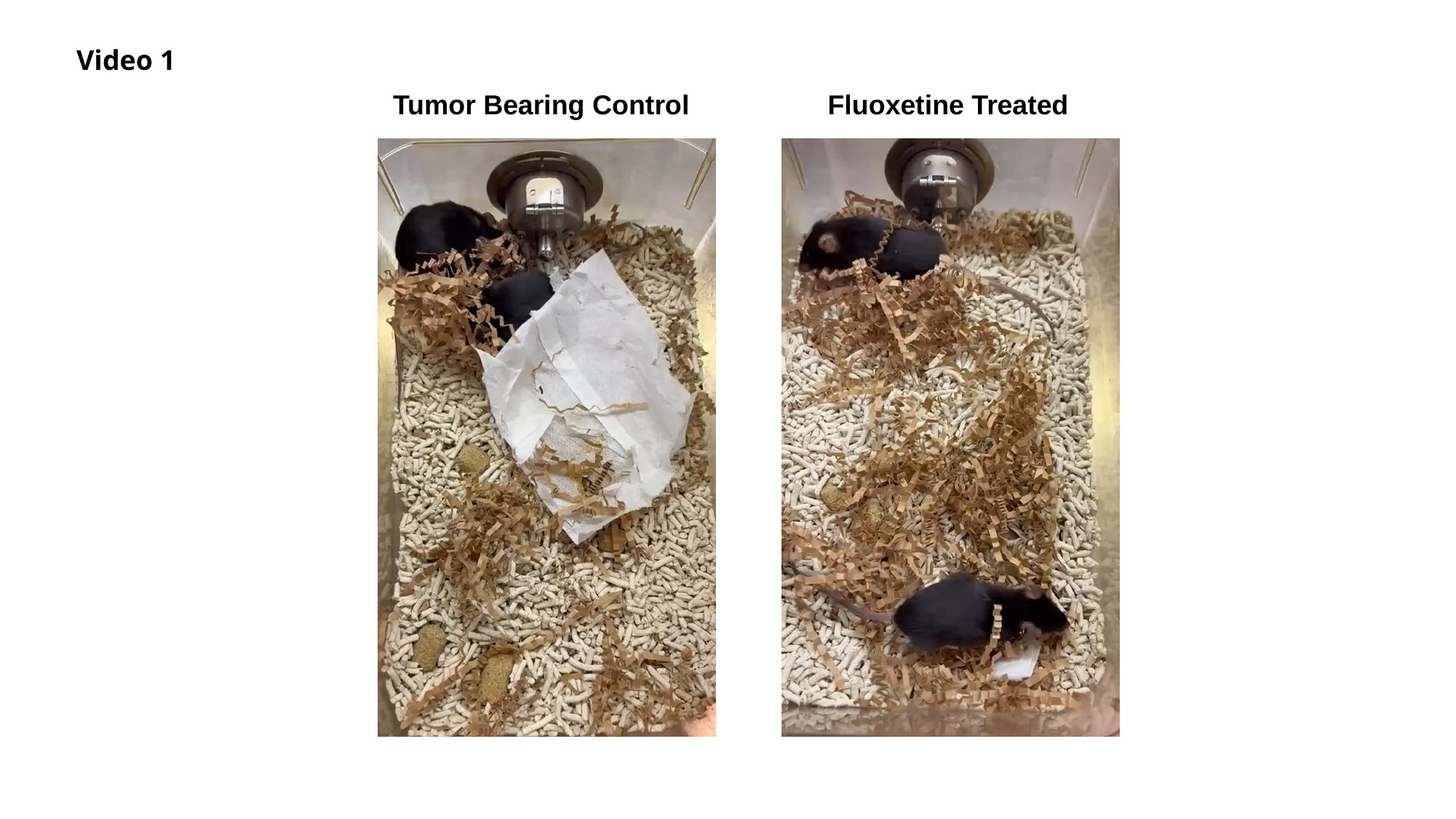

Video 1
Tumor Bearing Control
Fluoxetine Treated

### Slide 2
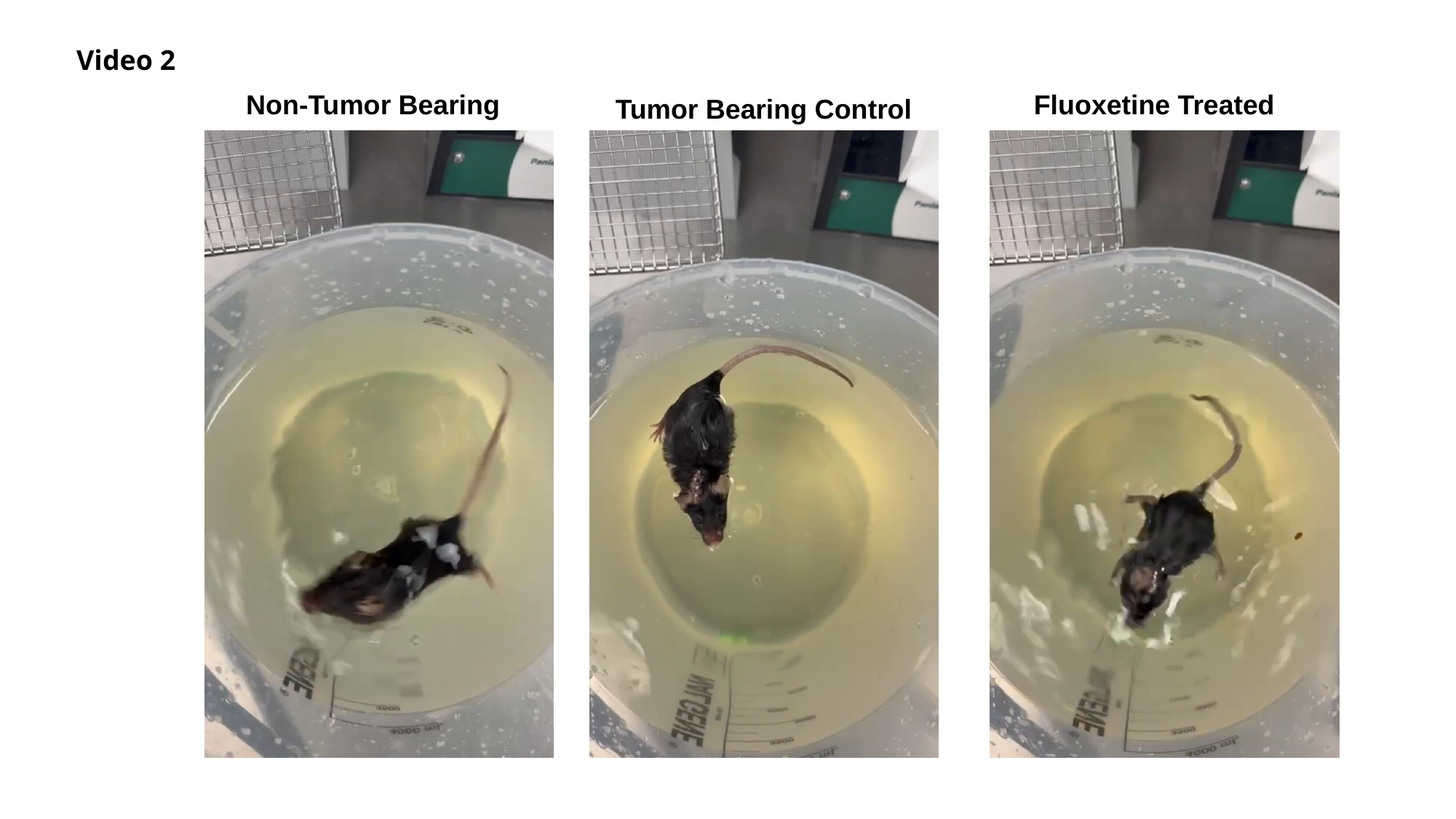

Video 2
Non-Tumor Bearing
Fluoxetine Treated
Tumor Bearing Control
